# Do microbial effects on hosts vary across life stage and vital rate?

**DOI:** 10.64898/2026.08.05.743066

**Authors:** Joshua C. Fowler, Gwendolyn B. Pohlmann, Eleanor M. Hay, Vicki W. Li, Zoe H. Meadors, Alma L. Reyes, Christopher A. Searcy, Michelle E. Afkhami

**Affiliations:** Department of Biology, University of Miami, Coral Gables, FL, 33146; Department of Environmental Studies, University of Colorado, Boulder, CO, 80303; Department of Ecology, Evolution, and Marine Biology, University of California Santa Barbara, Santa Barbara, CA, 93106

**Keywords:** symbiosis, microbial symbiont, mutualism, demography, life history, vital rates, life stage, meta-analysis

## Abstract

Interactions with microbial symbionts are widespread and can influence host fitness through effects on survival, growth, and reproduction. Demographic and evolutionary theory suggests that symbiont effects on components of host fitness should often vary in strength and even in direction as host and symbiont allocate limited resources. Through a meta-analysis of 1,736 host-symbiont effect size measurements, we evaluated how commonly the direction of symbiont effects varies across host life history stages and vital rates, and whether these effects across stages and vital rates differ across taxa. Contrary to our prediction, we found that, compared to a null expectation, published effects on host vital rates more often occurred consistently in the same direction, rather than in opposition. This finding suggests that symbiosis often provides benefits to host performance that manifest across vital rates. While the direction of symbiotic effect was often consistent, the magnitude of effects differed across fitness components and across host taxonomic groups, highlighting diverse demographic mechanisms by which symbiosis can influence host fitness. For hosts in the Poaceae family, we also found that effects of symbiosis varied across symbiont type: vertically-transmitted *Epichlöe* fungi most strongly benefited seed germination, while the benefits of mycorrhizal fungi and bacterial endophytes were stronger for adult growth and reproduction. Importantly, this literature survey also highlights gaps in the current literature on host-microbe symbiosis. Most studies measure growth during juvenile stages for taxa from iconic, well-studied symbioses. Meta-analytic estimates of symbiotic effects on host life history provide a path towards quantitatively modeling host-symbiont demography across the tree of life.

## Introduction

One of the earliest theoretical treatments of symbiosis noted the potential for symbionts’ effects on hosts to differ between multiple fitness components. Kostitzin (1934) observed that in some cases symbionts’ effects on the “vital coefficients” may be suitably captured with a single value, but that many cases require addressing the complication that symbionts’ effects on birth and death can vary and may depend on life stage/age. Over 90 years ago, Kostitzin bemoaned an “almost complete absence of quantitative data”, and that, “consequently, all attempts at theory are severely limited by the impossibility of experimental verification” (Kostitzin [1934] 1978). Today, there is extensive literature demonstrating the importance of symbiosis for host performance and evolution. For example, microbial symbionts have been implicated in protection from natural enemies in both insect (Oliver et al. 2003; Bruner-Montero and Jiggins 2023) and plant hosts (Panaccione et al. 2014), as well as enhancing host tolerance to drought (Decunta et al. 2021) and salinity stress (Subedi et al. 2022). The consequences of these effects scale up to influence populations (Yule et al. 2013; Fowler et al. 2024), species’ distributions (Afkhami et al. 2014; Fowler et al. 2023), ecosystem function (Rudgers et al. 2004; Alvarenga and Rousk 2022; Luo et al. 2023), as well as host evolution and diversification (Brucker and Bordenstein 2013; Petersen et al. 2023). Molecular sequencing has revealed that microbial symbionts are present within hosts across the entire tree of life (Rodriguez et al. 2009; Rosenberg et al. 2010), and describing how interactions vary across this diversity is a key step towards building general understanding of symbioses.

All organisms, both host and symbiont, must allocate their limited resources between growth, reproduction, or survival across their lifespans (Stearns 1989; Zera and Harshman 2001). Together, a host and its microbiota (collectively a holobiont) contribute to a shared phenotype, and it is this shared phenotype on which selection and evolution act (Guerrero et al. 2013; Simon et al. 2019). Therefore, symbionts can maximize their own fitness by investing their limited resources into aspects of their host’s life history that most benefit host fitness, and thus provide a higher return on investment to symbiont fitness. Many (but not all) microbial symbionts act to improve the resource acquisition ability of their hosts (e.g. mycorrhizal fungi that extend the capacity of their host plant to access both water (Ruiz-Lozano and Azcón 1995) and phosphorus (Bolan 1991)). Partnering with such a symbiont may alleviate resource limitation and allow their host to not only grow better, but also to be more likely to survive and reproduce throughout its lifetime than hosts without the partnership. Even symbioses that do not directly contribute to resource acquisition may indirectly increase resource availability and overall performance. For example, symbioses that mediate host growth through the production of hormones or pathogen control compounds, such as plant-growth promoting rhizobacteria (Glick 2012; Olanrewaju et al. 2017) or plant fungal endophytes (Waqas et al. 2012; Zhao et al. 2021; Mathew et al. 2023), could produce larger, healthier juveniles, which maintain growth advantages into adulthood. In turn, these larger hosts are likely to more successfully compete for and meet their resource needs (Plard et al. 2015). It has similarly been suggested that early life development of the human microbiome can have life-long impacts on immune responses (Martinez 2014) and chronic disease incidence (Sarkar et al. 2021) that contribute to overall health and performance.

At the same time, vital rates – defined as rates of survival, growth, and reproduction across different stages or ages – do not contribute equally to fitness (Pfister 1998; Caswell 2001). Thus, selection may target symbiont benefits toward the single vital rate with the greatest sensitivity. This suggests that microbial symbioses, many of which have long co-evolutionary histories with their hosts, may have strong fitness effects through manipulation of key vital rates, while their effects on low sensitivity vital rates may be negligible or even opposite in sign. For population fitness, the net interaction outcome depends on the magnitude of these effects multiplied by the sensitivity of each vital rate. For example, the grass *Cinna arundinacea* hosts an *Epichloë* fungal endophyte that reduced survival but increased reproductive output in a field trial (Rudgers et al. 2012). This *Epichloë* symbiont is vertically-transmitted (passed from maternal host to offspring), and the symbiont’s fitness is tightly linked to host fitness (Fine 1975). The overall result for *C. arundinacea* is a beneficial, mutualistic interaction, despite costs to survival, because host population growth has low sensitivity to changes in survival and thus there is little reward for the symbiont of *C. arundinacea* to invest in improving host survival. Focused investment by the symbiont to improve a single vital rate or life stage with high sensitivity sets the stage for an evolutionary trade-off, reducing investment in and potentially even negatively impacting other less sensitive fitness components. These sorts of conflicting effects have been documented across both animal (Martinez et al. 2015; Hoang et al. 2022) and plant (Gundel et al. 2006; Donald et al. 2021) hosts. The potential for conflicting effects of symbionts on hosts is supported by theoretical studies that have demonstrated that conflicting symbiont effects on fecundity and survival can serve as a mechanism underlying the evolution and stability of mutualisms (Ferriere et al. 2002; Travis et al. 2006; Fukui 2014). How frequently conflicting effects of symbiosis occur in nature is unknown.

Thus, we performed a meta-analysis to answer three central questions: (1) How does the magnitude of symbiotic effects on hosts differ across host vital rates and life history stages and do these patterns vary across host taxonomic groups? (2) how do symbiotic effects on hosts differ across symbiont type? And (3) how frequently are symbiotic effects in opposing directions across different life history stages and vital rates for a given host-symbiont pairing? Using the compiled dataset of 1,736 host-symbiont effect size measurements from experiments across the globe, we evaluated how the magnitude of effects varied across different life history stages, vital rates, and host families. Further, for Poaceae, the host family that associated with the widest diversity of symbiont taxa, we tested how the magnitude of effects varied across symbiont types, including vertically-versus horizontally-transmitted fungal and bacterial symbionts. Overall, we expected that opposing vital rate effects would be common because symbionts have limited resources to allocate towards influencing host fitness. However, many symbioses also expand the available resource pool for their hosts, which could allow these partners to benefit multiple fitness components simultaneously. While this work does not directly evaluate these underlying resource allocation mechanisms, we lay the groundwork for future studies of the demography of host-microbe symbiosis. By compiling studies that quantify symbiotic effects on host vital rates across a broad suite of host-microbe symbiota and life history stages, our analysis fills the long-standing call for better experimental estimates of the mean and variance in mutualist effects on hosts’ “vital coefficients” while also identifying any taxonomic gaps in the systems in which these demographic effects of symbiosis have been investigated.

## Methods

### Data Collection

To collate estimates of symbionts’ effects on host vital rates, we performed a systematic literature search in Web of Science (Siddaway et al. 2019). On Feb. 21, 2024, we collected references identified through the search term: “ALL=((symbio*) AND ((microb* OR fung* OR bacteri* OR alga*) OR (mycorrhiz* OR rhizobi* OR endophyt* OR ectomyc* OR endomyc* OR arbuscul* OR wolbach* OR epichlo* OR buchner* OR symbiodini* OR burkholder* OR entomocortici* OR vibri* OR endozoicomon*)) AND ((fitness OR performance OR vital rate OR demograph* OR population growth) OR (survival OR biomass OR growth OR fecundity OR germination OR hatch)) AND (experimen* OR inoculat* OR steriliz*))”. This search resulted in 9,606 articles. During our screening of articles, we included studies that (i) provided estimates of host performance for both symbiotic and aposymbiotic hosts, (ii) included treatments that focused on the presence/absence of one host-microbe pairing at a time, which included both experimental manipulations and comparisons of naturally occurring symbiotic and symbiont-free individuals, and (iii) provided measurements of the mean and dispersion (s.d. or s.e.) around the mean of host vital rates across these treatments. After examining the first 200 search results sorted by relevance to the search terms, 78 articles met our criteria (Fig. S14, Fig. S15). We repeated our search of Web of Science to account for regional spellings variations (i.e., sterilisation vs. sterilization) and found that the studies included in the top 200 most relevant search results were unchanged. While we did not seek to exclusively search for beneficial symbioses as our questions related to how symbiotic effects can be both positive or negative for a given interaction, our search terms included several well-known, commonly mutualistic symbioses.

Across these 78 articles, 106 unique experiments provided 1,736 measurements of symbiotic effects on host vital rates. For each measurement, we documented the focal vital rate (growth, reproduction, survival, or recruitment (i.e. processes that add new individuals to populations, such as germination or hatching)) as well as the host life stage (embryo, juvenile, adult). Here, we use the term “embryo” to refer to organisms in seed or larval categories, which are typically involved in recruitment, and before typical juvenile structures emerge (e.g., the first cotyledons emerging from seeds). We defined organisms as “juvenile” for any measurement before reproductive maturity unless otherwise indicated by study authors. If reproductive timing was not specified or clear from the biology of the study organism (e.g., long-lived tree species), we defined individuals older than one year as “adults”, however we note that the majority of studies in the dataset were conducted for less than one year. In some cases, articles provided multiple temporally offset measurements of fitness effects within a single life stage (e.g., measurements at different points across juvenile development) and for these we recorded juvenile stages 1,2,3…n. Shorter term measurements of individual fitness components during a particular stage may not encompass the complete life stages, but still provide information on the rate of e.g., survival or germination during different life stages. We extracted data on the mean, standard deviation, and sample size of each fitness component either directly from the text or from published plots using the metaDigitize package in R (Pick et al. 2019). For each study, we also recorded the experimental context (lab, greenhouse, field) and whether the symbiotic treatment was experimental or observational (i.e., whether the hosts were naturally symbiotic or inoculated, or whether hosts were naturally symbiont free or disinfected). Analysis of differences across experimental contexts and symbiont treatment types is provided in the Supplemental Methods, and a map of the study locations is provided in the Supplementary Figures (Fig. S13).

### Effect size calculation

We calculated a standardized effect size, the Relative Interaction Intensity (RII) following Armas et al. 2004 for each measurement.

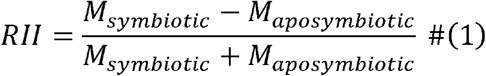

Where M_symbiotic_ are M_aposymbiotic_ are the mean values of measurements with and without symbionts respectively. Each RII value represents a measurement of symbiont effects on a vital rate during a particular life history stage (e.g., positive or negative effect of partnering with a symbiont on germination rate). RII is bounded between 1 and −1 and is symmetrical around zero. These properties allow for comparison between both negative and positive effects of symbiosis on host vital rates on the same scale and are also particularly useful for obligate effects of symbiosis (e.g., 0% germination in absence of symbiont), for which other effect sizes, such as the log response ratio, would be undefined (Armas et al. 2004; Chamberlain et al. 2014).

### Describing taxonomic gaps in studies of host-microbe symbioses

We first described the taxonomic diversity of hosts included in the dataset, then visualized the frequency of measurements across different vital rates and life stages to identify potential gaps in knowledge. To connect recorded host and symbiont taxa to taxonomic information, we used the R package *rotl* (Michonneau et al. 2016). We additionally evaluated the extent to which small-study bias (the potential for low-precision studies to publish higher than expected effect sizes) may have shaped measured effect sizes within the compiled dataset. In brief, we performed a regression analysis analogous to an Egger’s test (Egger et al. 1997; Schwarzer et al. 2015), implemented in a Bayesian framework (Nakagawa et al. 2022). An Egger’s test evaluates the relationship between scaled effect sizes and study precision, where an intercept different from zero suggests potential publication bias. Details of this analysis are provided in the Supplemental Methods.

### Evaluating variation in symbiont effects across vital rates and life stages

We used a Bayesian meta-regression approach to explore variation in published symbiotic effects on hosts, and how these effects vary across taxa. To quantify how symbiont effects varied across vital rates and life stages, we modeled observed effect sizes, RII, as a

Gaussian distribution with measurement error. We built separate models to analyze differences across vital rates and across life stages because some vital rate and life stage combinations were inherently conflated (e.g., reproduction only occurred during adult life stages). Both models shared the same overall structure, described by Eqn. 2.

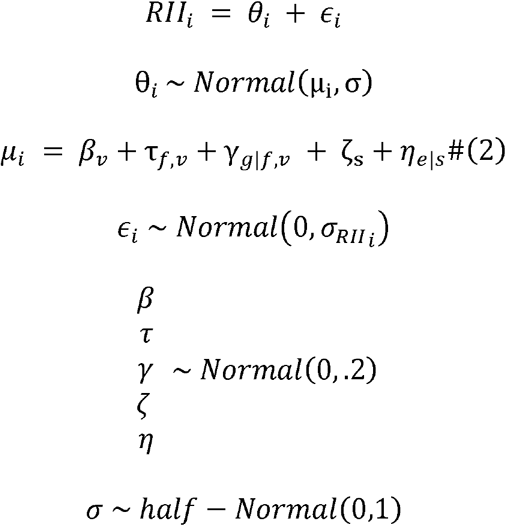

Here, θ is a distribution of true effect sizes for each estimate (*i*) with error ɛ. We modeled heterogeneity in effect sizes with intercepts that vary according to vital rate descriptor (v) (i.e. either different vital rates (survival, growth, reproduction or recruitment) or different life stages (embryo, juvenile, adult)). We incorporated nested random effects describing host taxonomy τ accounting for host family (*f*), and γ accounting for host genus (*g*) that were allowed to vary by vital rate component. We additionally included study-level random effects (*η*and ζ) accounting for multiple separate measurements of effect size from each experiment (e) within each paper (s) from which we extracted data. Finally, the across study estimate of measurement error (ɛ) depended upon the measured standard deviation in RII σ_RII_ which we calculated following Armas et al. 2004. All parameters were given weakly informative priors (Gelman et al. 2017). For β, τ, γ, ζ, and*η*, we first tested normally distributed priors with mean = 0 and sd = 1, and placed a positively defined half-normal distribution with mode = 0 and sd = 1 on the variance term σ. This set of prior choices resulted in small numbers of divergent Markov chain Monte Carlo (MCMC) transitions during model fitting. We re-fit the model with narrower priors for the linear predictor parameters (Gaussian distribution with mean = 0 and sd = 0.2) that constrained most estimates to the scale of RII (-1 to +1). Narrower priors facilitated model convergence and provided qualitatively similar model fits to alternatives with wider priors. While we considered alternative models that explicitly incorporated truncation bounds, sparse data across combinations of fitness components and taxonomic groups made estimating truncation bounds challenging. The Gaussian model adequately captured group-level means in posterior predictive checks (Figs. S9-S10), allowing inference on differences in average effect size across fitness components and taxonomic groups.

### Evaluating how symbiont type influences symbiont effects across vital rates and life stages

We next explored how symbiotic effects on host performance metrics differed across different symbiont partners. Using data from the grass family (Poaceae), which associated with the broadest diversity of symbiont taxa, we modeled observed effect sizes, RII, with a meta-regression as described above, modified to incorporate parameters that allow estimates of RII to vary according to type of symbiont. We categorized the symbionts as either *Epichloë* fungi (predominantly vertically-transmitted, above ground endophytes), mycorrhizal fungi (below ground, root associations), or bacteria (including both above and belowground symbionts) (Table S1. Eqn. 3 describes the linear predictor:

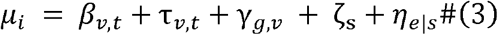

where the intercept and slope parameters (βand τ) vary according to symbiont type (t) in addition to vital rate descriptor (v). Because this analysis focused on a single host family, note that this model did not include a family-level random effect, but the rest of the model structure and priors remained as described above in Eqn. 2.

We performed all statistical analyses using the *brms* package in R (Bürkner 2017). Each model was sampled for 1000 warmup and 1000 sampling iterations across four MCMC chains. We evaluated model convergence by ascertaining that R̂ values were below 1.01 (Gelman and Rubin 1992; Vehtari et al. 2021), and visually assessed model fit with graphical posterior predictive checks (Figs. S9-S10) (Gelman et al. 1996).

### Evaluating how frequently symbiotic effects are in opposing directions

To evaluate how frequently symbiotic effects on vital rates were in opposing directions, we performed a permutation test. For each unique host-microbe pairing within the meta-analytic dataset with more than one measurement of a symbiotic effect across different vital rates or life stages (n = 199 unique symbiota), we determined whether symbiotic effects were consistently positive, consistently negative, or included at least one effect in an opposing direction. For a given unique host-symbiont pairing, we calculated the proportion of all symbiotic effect estimates that were in the same or opposing direction. We then compared the frequency of these outcomes with 500 random permutations of the data. This allowed us to evaluate, given the relative proportion of positive and negative symbiotic effects present in our dataset, how commonly one should expect opposing vital rate effects to occur by chance. We repeated this analysis sub-setting the data for each host family to evaluate if particular taxa differ from the overall trend.

## Results

### Is there evidence of publication bias or data limitation within the dataset?

Measured effects of microbial symbionts on hosts came from 240 unique host symbiont pairings from 59 host genera, 27 families, and 22 orders (Fig. 1), interacting with symbionts from 13 families of bacteria, 15 families of fungi, and one genus of algal symbiont. Among 59 genera of hosts, 51 were plants (n = 1,668 effect size measurements), 7 were arthropods (n = 58 effect size measurements), and one was a coral host (n = 10 effect size measurements). The most commonly measured interaction types were between grasses (Poaceae) and their fungal symbionts (576/1736 effect size measurements (33%)), including both above-ground endophytes in the genus *Epichloë* (n = 14 unique host symbiont pairings), mycorrhizal fungi (n = 12 unique host symbiont pairings), and bacterial symbionts (n = 8 unique host symbiont pairings). The next most common were interactions between the orchids (Orchidaceae) and their specialized fungal symbionts (393/1736 measurements (23%)). The average number of effect size measurements per host-symbiont pairing was 4.2 across combinations of vital rate metrics and life stages, while 41 pairings had measurements for only one metric. The host-symbiont pairing with the most distinct measurements was *Catalpa bungei* (a tree native to China in the family Bignoniaceae) and its interaction with the arbuscular mycorrhizal fungus *Rhizophagus intraradices*, with 33 measurements across two life stages for a diversity of vital rate metrics. It is worth noting that while some taxa had multiple vital rate metric measurements, these were concentrated in only one or a few metric categories. For example, measurements for hosts within the Bignonaceae and Lamiaceae families provided relatively high proportions of measurements (63 and 37 measurements, respectively), but these were entirely focused on symbiont effects on host growth (including different dimensions of growth such as biomass, height, or branching number), with no estimates of effects on survival, reproduction, or recruitment for these taxa (Fig. 1A).

**Figure 1.**
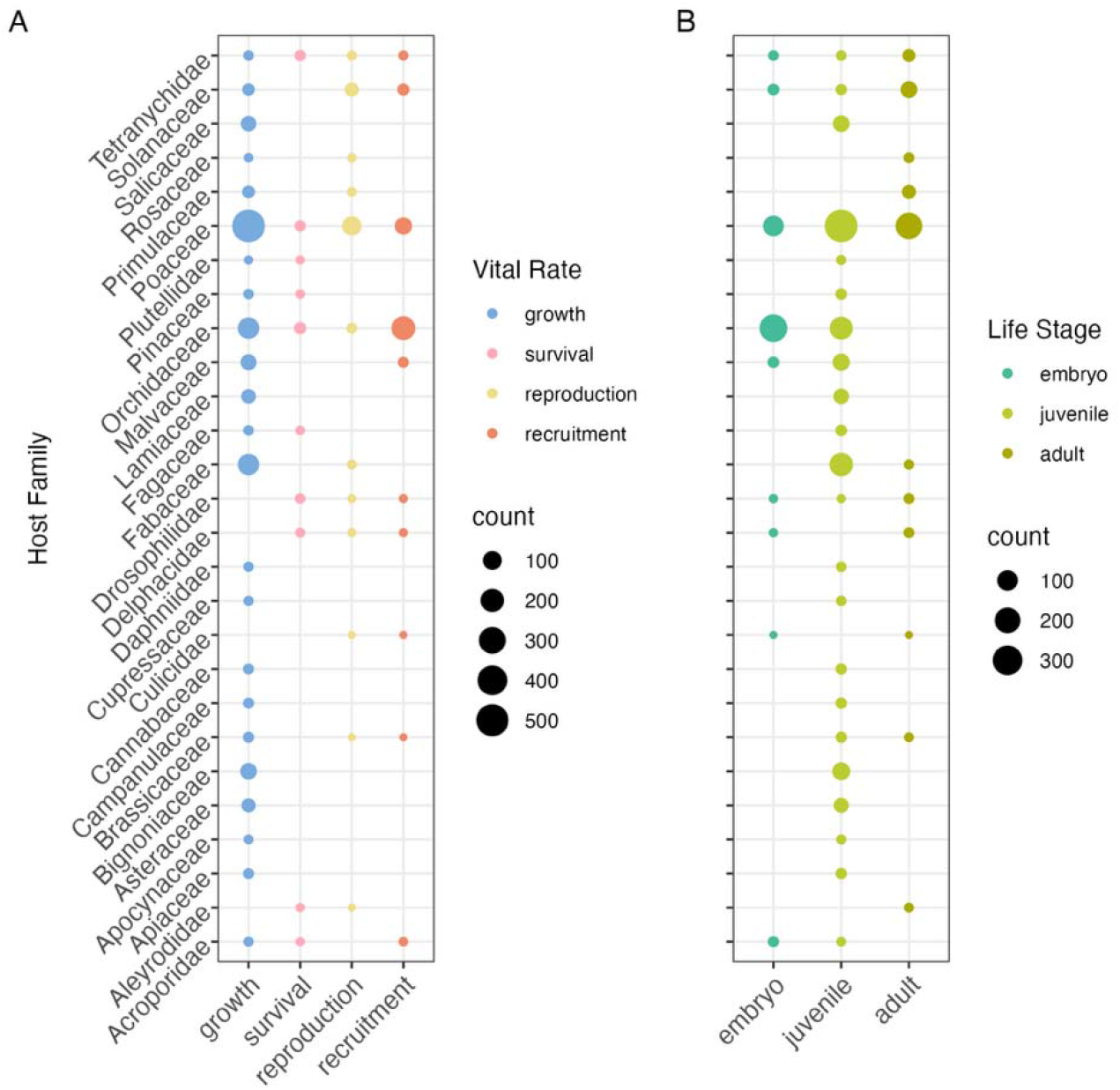
Counts of host-symbiont demographic effects across taxonomic diversity. (A) Counts of effect sizes according to host vital rate and host taxonomic family. (B) Counts of effect sizes according to host life stage and host taxonomic family. Note the prevalence of juvenile growth rate measurements. Point size reflects the count of measured effect sizes, and colors represent the measured vital rate (blue: growth, pink: survival, yellow: reproduction, orange: recruitment) or life stage (turquoise: embryo, lime: juvenile, olive: adult).

Across the dataset, the most common measurements were of host growth (1176/1736 measurements (68%)) (Fig. 1A) and were taken during juvenile life stages (1008/1736 measurements (58%)) (Fig. 1B). Recruitment was the next most commonly measured demographic metric with 329 measurements (19%), followed by reproduction (174; 10%), and then survival (57; 3.3%). There were 391 measurements of the embryo life stage (22.5%) and 337 measurements of adults (19.5%). The large majority of symbiota were measured under controlled conditions such as in greenhouses or growth chambers (1456/1736 measurements; (85%)) rather than in field experiments (260/1736 measurements (15%)). After accounting for host taxonomy and measurement error (see Supplemental Methods), we did not find differences in the magnitude of symbiont effects between experimental manipulations of symbiont presence/absence vs. naturally occurring variation or between field vs. greenhouse vs. lab settings (Fig. S4 – S8).

We did not find evidence that publication bias contributed to significant variation in the compiled dataset. Full details of our analysis to test for potential publication bias are provided in the Supplemental Methods. In brief, based on an Egger’s test (Egger et al. 1997; Schwarzer et al. 2015), we found that 95% posterior credible intervals for regression intercepts did not differ from zero, indicating no statistical evidence of “small study” publication bias (Fig. S1B; Fig. S2B), a result that held across each vital rate and life stage.

### How do microbial effects vary across host vital rates?

Measurements of symbiotic effects spanned from completely beneficial (RII = 1) to completely antagonistic (RII =-1) for hosts, but on average, effects were consistently positive across vital rates (Fig. 2A). Averaged across variation associated with host taxonomy and study heterogeneity, the mean RII was positive for all four vital rates. The magnitude of effects on reproduction was greatest (*mean* = 0.19, 95% *CI* = [0.06, 0.31]), followed by marginally weaker effects on growth (*mean* = 0.16, *CI* = [0.07, 0.25]), and then survival (*mean* = 0.11, *CI* = [-0.12, 0.33]) and recruitment (*mean* = 0.10, *CI* = [-0.06, 0.24]) (Fig 2A). Each of these metrics also had high certainty that true effects were greater than zero; growth and reproduction had greater than 95% posterior probabilities of positive effects, while recruitment and survival had greater than 87% and 83% posterior probabilities of positive effects, respectively.

**Figure 2.**
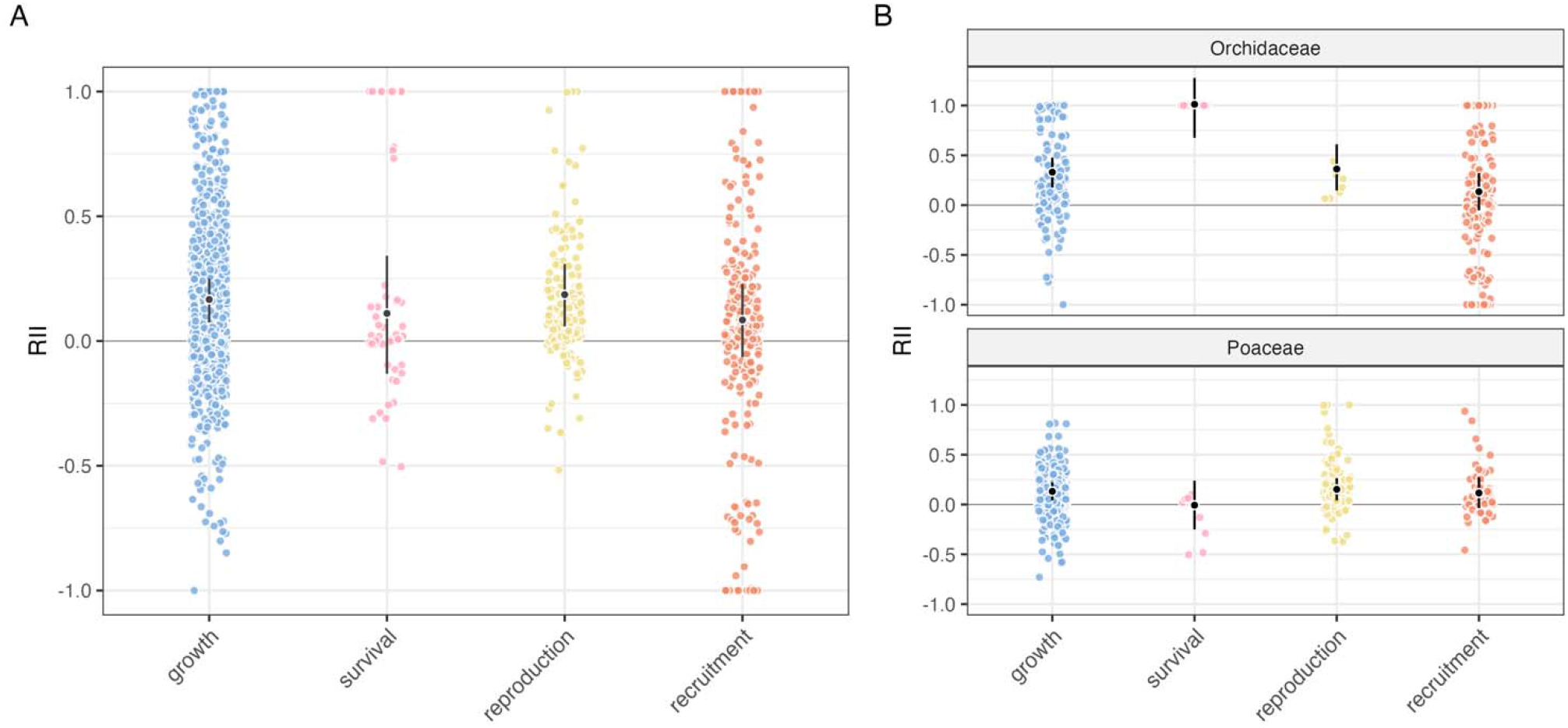
Symbiotic effects are consistently positive on average across vital rates. Meta-analytic means (black) along with 95% CI are shown for each vital rate type (growth, survival, reproduction, and recruitment), along with observed RII values across studies (colored points). (A) Effect sizes averaging across taxonomic diversity are shown along with (B) effect sizes for the two most commonly recorded host families within the dataset, Orchidaceae and Poaceae, highlighting taxonomic variation in symbiotic effects.

Evaluating model predictions across the most commonly measured families within the dataset showed that symbiotic vital rate effects were positive across taxa. However, we also found interesting variation. For example, when examining the two most recorded host families in our dataset (Orchidaceae and Poaceae), the relative impact of symbiotic benefits differed per vital rate (Fig. 2B). Within Orchidaceae hosts, which included many highly obligate orchid-mycorrhizal symbioses, symbionts most strongly affected survival (*mean* = 1.0, *CI* = [0.68, 1.31]; >99% positive posterior probability), while for Poaceae, symbionts most strongly affected growth (*mean* = 0.13, *CI* = [0.05, 0.22]; >99% positive posterior probability) or reproduction (*mean* = 0.15, *CI* = [0.05, 0.26]; >95% positive posterior probability). Our analysis indicated a greater than 99% posterior probability that symbiotic effects on survival for Orchidaceae hosts were stronger than those for Poaceae hosts, and effects on Poaceae survival were neglible (*mean* =-0.001, *CI* = [-0.25, 0.26]; 47.8% positive posterior probability). While some families showed strong effects on particular vital rates – such as Orchidaceae and Pinaceae that experience strong positive survival effects (Fig. S11) – effects of symbionts were modest and consistent across vital rates for many taxa. For example, within the Tetranychidae hosts, a clade of spider mites, all the estimated symbiotic effects were weakly positive to a similar degree across vital rates (growth = 0.13, *CI* = [-0.09, 0.33], (90% positive probability); reproduction = 0.14, *CI* = [-0.12, 0.36], (87% positive probability); recruitment = 0.06, *CI* = [-0.24, 0.32], (67% positive probability); survival = 0.10, *CI* = [-0.26, 0.43], (74% positive probability)), with posterior credible intervals that each overlapped zero (Fig. S11).

### How do microbial effects vary across host life history stages?

Symbiotic effects on hosts were consistently positive regardless of host life history stage (Fig. 3A). Averaging across taxonomic diversity, we found strong positive symbiotic effects on juvenile (*mean* = 0.19, *CI* = [0.08, 0.30]) and adult (*mean* = 0.11, *CI* = [0.02, 0.19]) stages, each of which had greater than 99% posterior probability of positive RII. Our analysis indicated relatively weaker positive effects of microbial symbiosis with wide variation for embryonic hosts (*mean* = 0.08, *CI* = [-0.06, 0.23]). On average, effects on embryonic life stages had an 87% posterior probability of positive RII values (Fig. 3A). Effects on juvenile stages were 90% more likely to be greater than effects on embryonic or adult life stages.

**Figure 3.**
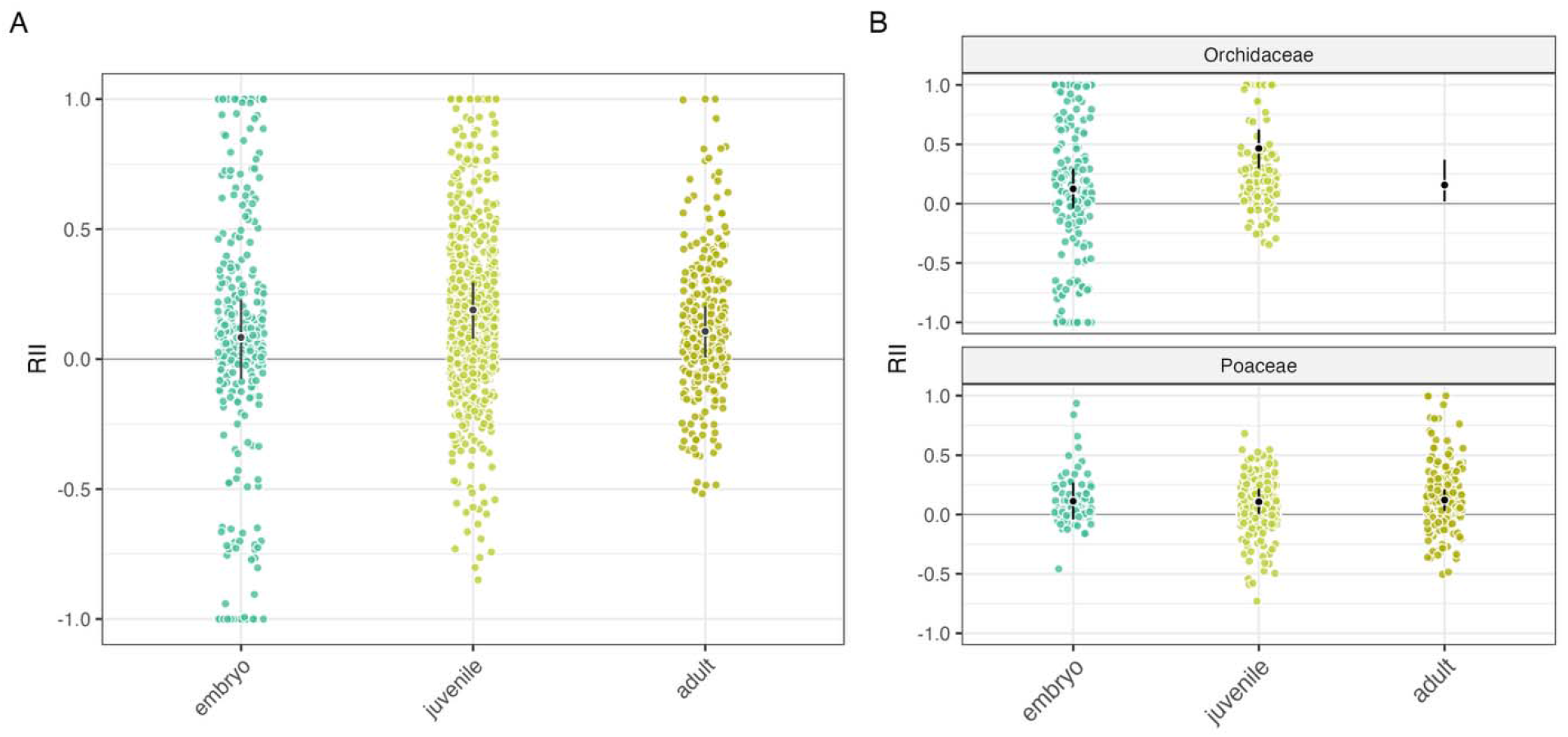
Symbiotic effects are consistently positive on average across host life stages. Meta-analytic means (black) along with 95% CI are shown for each life stage category (embryo, juvenile, or adult), along with observed RII values across studies (colored points). (A) Predictions averaging across taxonomic diversity are shown along with (B) predictions for the two most commonly recorded taxonomic families within the dataset, Orchidaceae and Poaceae.

We also evaluated how these effects varied across host taxa and found that, within the most commonly measured families, symbiont effects on embryo stages were weak relative to juvenile and adult stages (Fig. 3B). In Poaceae, both effects on juvenile stages (*mean* = 0.11, *CI* = [0.01, 0.21]) and effects on adults (*mean* = 0.12, *CI* = [0.03, 0.21]) had greater than 97% posterior probability of positive effects, while effects on embryonic stages (*mean* = 0.12, *CI* = [-0.03, 0.27]) had an 93% positive posterior probability. For Orchidaceae, there was clear evidence that symbiont effects were strongest on juvenile stages relative to embryonic stages, while there were no measurements for adult stages. In Orchidaceae, effects on juvenile stages (*mean* = 0.46; *CI* = [0.27, 0.60]), had greater than 97% posterior probability of positive effects, while effects on embryonic stages (*mean* = 0.12, *CI* = [-0.04, 0.31]) had a 92% positive posterior probability.

There was a greater than 97% posterior probability that effects on juveniles were more positive than effects on embryonic or adult life stages. There was also clear evidence that microbial effects on juvenile stages were particularly strong in Orchidaceae relative to other measured host families (Fig. S12). We found a 95% posterior probability that symbiotic effects on juvenile Orchidaceae hosts were greater than effects on juveniles across all other host families. There were no measurements of symbiotic effects among adult stages in Orchidaceae, likely due to the inability of the host to live to later life stages without its obligate mycorrhizal symbionts, and estimated effects on adult stages were influenced by partial pooling in the hierarchical model (*mean* = 0.21, *CI* = [0.01, 0.36]).

### How do symbiotic effects on hosts differ across symbiont type?

In this analysis within Poaceae hosts, we found that the magnitude of symbiotic effects on particular vital rates and life stages differed across symbiont types. Averaging across host genera, endophytic *Epichloë* fungi, which are commonly vertically-transmitted through seeds, provided clear positive effects on recruitment (*mean* = 0.20, *CI* = [0.03, 0.38]; 98% positive posterior probability), weaker positive effects on reproduction (*mean* = 0.13, *CI* = [-0.03, 0.28]; 95% positive posterior probability) and growth (*mean* = 0.08, *CI* = [-0.03, 0.18]; 93% positive posterior probability), and negligible effects on survival (*mean* =-0.01, *CI* = [-0.22, 0.18]; 44% positive posterior probability) (Fig. 4A). In contrast, belowground mycorrhizal fungi had negligible or weakly negative effects on recruitment (*mean* =-0.05, *CI* = [-0.24, 0.16]; 30% positive posterior probability) (Fig. 4A), while their strongest positive effects were on growth (*mean* = 0.11, *CI* = [-0.01, 0.22]; 97% positive posterior probability). Effects on reproduction for mycorrhizae were weaker (*mean* = 0.10, *CI* = [-0.10, 0.30]; 85% positive posterior probability), and there were no measurements of effects on survival. Across bacterial symbionts, there were strong positive effects on reproduction (*mean* = 0.22, *CI* = [0.03, 0.41]), growth (*mean* = 0.14, *CI* = [0.04, 0.25]), and recruitment (mean = 0.12, *CI* = [-0.04, 0.27]), which had 99%, 99%, and 94% positive posterior probabilities, respectively. The dataset did not include measurements for bacterial symbiont effects on survival.

**Figure 4.**
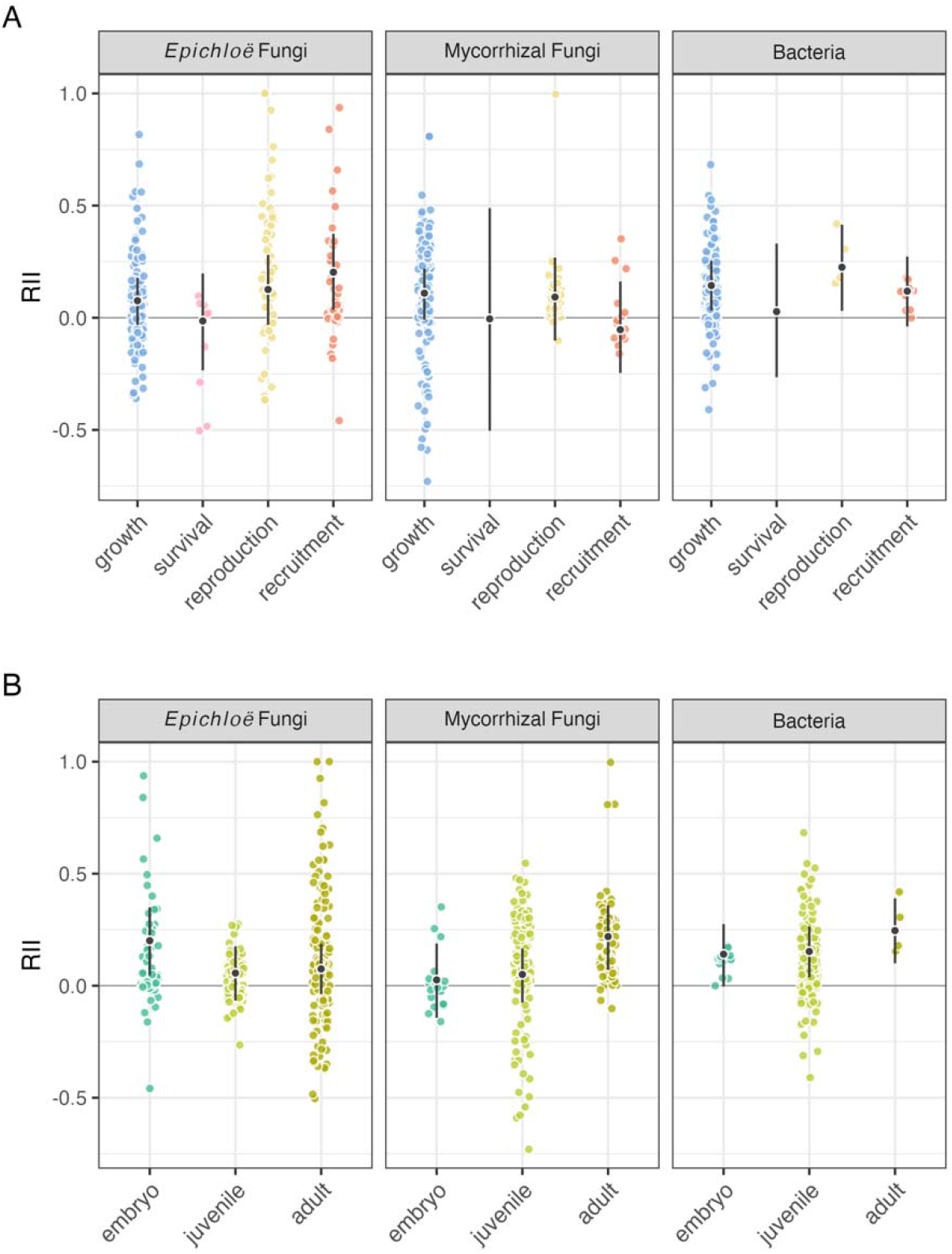
Symbiotic effects on host vital rate and life stage depend on symbiont identity. Vertically-transmitted Epichloë fungal endophytes have their strongest effects on embryos/recruitment, while horizontally-transmitted symbionts have their strongest effects on adult growth and reproduction. Meta-analytic means (black) along with 95% CI are shown (A) for each vital rate type along with colored points representing observed RII values across studies (blue: growth, pink: survival, yellow: reproduction, or orange: recruitment), and (B) for each life stage category (turquoise: embryo, lime: juvenile, olive: adult).

Symbiotic effects across life stages also differed for each symbiont type. For example, the strongest effects of *Epichloë* symbiosis were during embryonic life stages (*mean* = 0.20, *CI* = [0.04, 0.36]; 99% posterior probability of positive effects). There was a 96% probability that effects on embryonic host stages were greater than on other host life stages for *Epichloë* symbionts. In contrast, the effects of mycorrhizal (*mean* = 0.21, *CI* = [0.08, 0.36]) and bacterial (*mean* = 0.25, *CI* = [0.10, 0.39]) symbionts were strongest during adult life stages (both 99% posterior probability of positive effects). For these symbionts, there was greater than 99% and 91% posterior probability that effects on host adult stages were greater than other host life stages. Effects of symbioses were generally positive across life stages, with posterior probabilities ranging from 79% to >99% positive, with the exception of mycorrhizal effects at the embryo stage, which were negligible (*mean* = 0.03, *CI* = [-0.13, 0.18]; 65% positive posterior probability).

### How common are opposing vital rate effects?

Finally, we evaluated the frequency at which symbiotic effects that are opposite in sign occur for a given host-microbe pairing. We found that the observed dataset contained a higher proportion of consistently positive vital rate effects than expected by chance (Fig. 5). We found 79 out of 199 symbiota had only positive measurements of symbiotic effects reported, while the expected mean of the permutated datasets was 50.9 symbiota. The observed value was outside of the 95% CI of permuted datasets (41 - 61 symbiota). Symbiota that had opposing measurements were less frequent than expected (105 vs 145.9 expected), which was also outside of the 95% CIs of permuted datasets (136 – 156 symbiota). Those that had only negative measurements were more frequent (15 vs 2.1 expected) compared to the permuted data, which was also above the 95% CI of the permuted datasets (0 – 5 symbiota).

**Figure 5.**
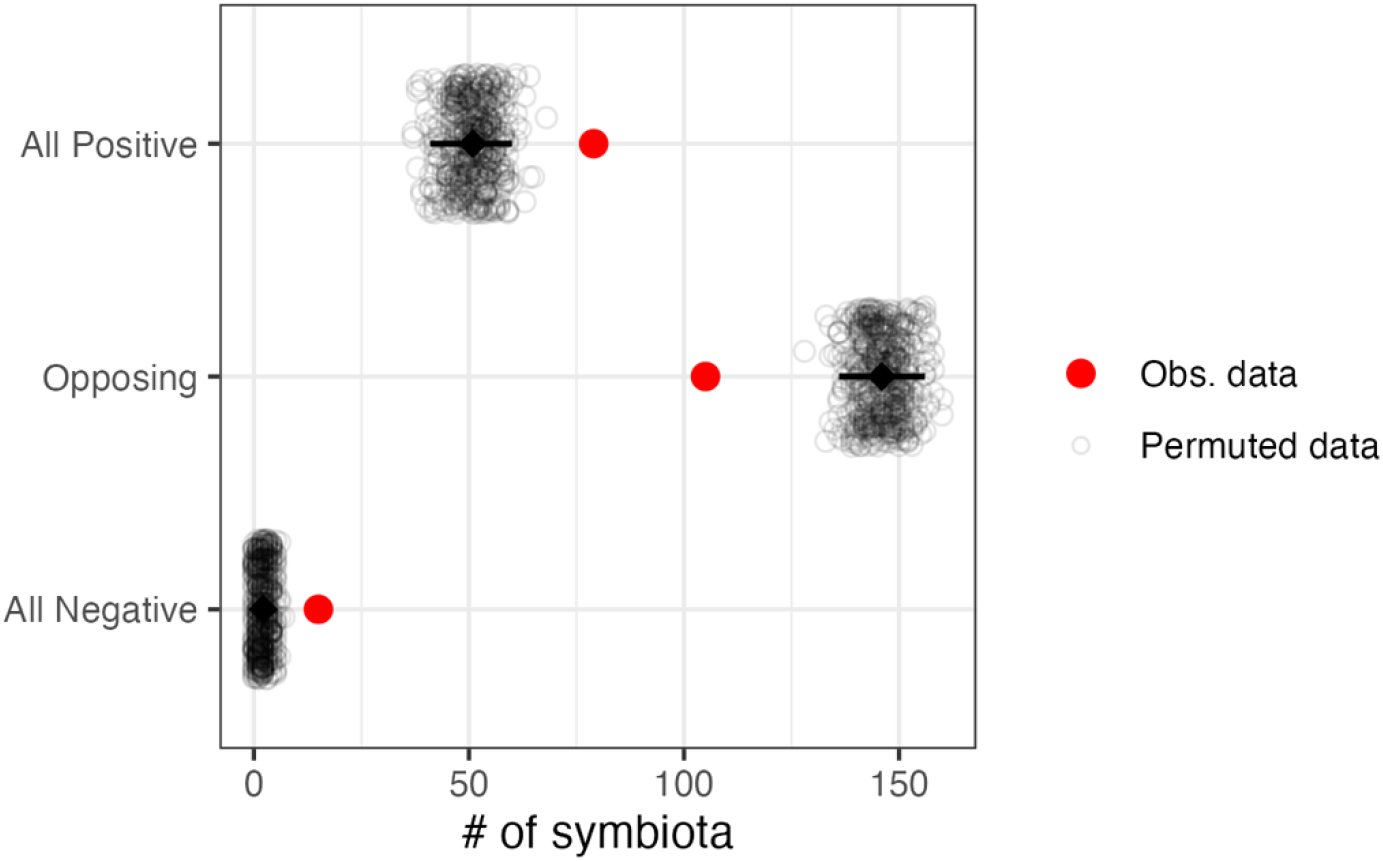
Observed symbiotic effects were less commonly in opposing directions than expected by chance. Points represent the number of unique host-microbe symbiota with either all positive vital rate effects, at least one opposing vital rate effect, or all negative vital rate effects within the observed meta-analytic dataset (red) and across 500 randomly permuted datasets (black). Black diamonds represent the expected mean for each category from the permuted datasets along with error bars extending along the 95% CI.

Repeating this permutation analysis for each host family with more than one unique host-symbiont pairing showed that high frequencies of consistent positive effects and low frequencies of opposing symbiotic effects were common across families (Fig. S13). The Orchidaceae family contributed the majority of the consistently positive effects, and surprisingly, also the majority of the consistently negative effects (Fig. S13). There were no families that had opposing symbiotic effects greater than expected by chance. However, some families such as the Malvaceae and Lamiaceae did have frequencies of opposing symbiotic effects within the 95% CI of the permuted datasets (Fig. S13). Families represented in the dataset by only a few unique symbiota did not differ from expected frequencies, but for several of these families, only positive effects were documented (Fig. S13).

## Discussion

Variation in symbiotic effects across different components of host life history has the potential to drastically alter consequences for host populations and communities. We originally predicted that opposing vital rate effects would be common because selection would favor beneficial symbionts to invest limited resources in manipulating vital rates that make particularly high contributions to fitness. In contrast, we found that symbiotic effects were consistently in the same direction (either all positive or all negative). Consistently positive symbiont effects were the most commonly observed outcome across host vital rates, life stages, and taxa. This contrasts with evolutionary theory suggesting that conflicting effects contributes to the establishment and stability of mutualism (Ferriere et al. 2002; Travis et al. 2006; Fukui 2014). The lack of opposing effects suggests that, in part, the benefits of these interactions come from general benefits across life history processes that we speculate could be the result of symbionts that expand the available resource pool for their hosts. This could occur directly (such as via improved acquisition of multiple resources by mycorrhizal fungi (Ruiz-Lozano and Azcón 1995; Bolan 1991)), or through indirect effects across life history processes (improved host growth during early life stages also benefits later host survival, reproduction, or the quality of future offspring) (Angilletta Jr. et al. 2006; Plard et al. 2015).

Our meta-analysis of 1,736 measurements of symbionts’ effects on host vital rates also revealed meaningful variation in the magnitude of symbiotic effects across different vital rates, life stages, and taxa. Despite the fact that opposing effects were uncommon, for some taxa, symbiotic effects were strongest on particular metrics or stages, while effects on other metrics were weaker. This was most evident in the survival of juvenile stages for orchid-mycorrhizae symbioses. While our analysis was agnostic to vital rate sensitivities given the scarcity of demographic models for these host species, it is likely that these highly specialized symbioses invest in what are commonly higher sensitivity life history processes: survival and growth rather than reproduction (Pfister 1998), and the success of established juveniles rather than high variance seed germination (Crouse et al. 1987; Sæther et al. 2013). In contrast to Orchidaceae, we found that several other host families exhibited strong symbiotic effects on reproduction. For example, the symbionts of Poaceae hosts positively influenced reproduction, growth, and recruitment, while having negligible or negative effects on survival. Among these symbionts, highly specialized *Epichloë* fungal endophytes (which are predominantly, but not exclusively vertically transmitted) had strong positive effects on recruitment and success of seeds. This contrasts with the effects of other fungal and bacterial symbionts of grasses that are horizontally acquired from the soil with each generation. These horizontally-transmitted symbionts had negligible effects on recruitment, but strong positive effects on adult growth and reproduction. This highlights that the optimal investment strategy for some symbioses may be to maximize host reproductive success rather than survival, depending on both how these processes contribute to host fitness and on symbiont life history traits (e.g., lifespan within the host, reproductive rate, dispersal ability). The picture may even be more complicated when considering the sequential events across a host’s lifespan that contribute to successful reproduction and, for the symbiont, successful transmission (Stott et al. 2024). While we did not find opposing vital rate effects to be common, we would expect that symbiotic effects that target highly sensitive fitness components may be common, especially for highly specialized and obligate interactions.

Fully understanding how symbionts contribute to population growth requires quantifying effects on each stage of host life cycles and weighting these symbiont effects by the sensitivity of population growth to that vital rate or stage in the context of a demographic analysis (Yule et al. 2013). From a practical standpoint, the result that symbiotic effects typically occur in the same direction is good news because it means that studies, which commonly measure one or only a few metrics of host performance, are likely to correctly predict the direction of other unmeasured demographic effects. However, studies that only measure one or a few vital rate metrics likely underestimate the impact that the symbiosis truly has for population growth because correlated contributions occur on other vital rates across the host life cycle. Using approaches that leverage information across datasets (Schaub and Kéry 2012; Chevalier and Knape 2020), as we have done through this meta-analysis, is one potential path towards estimating the effects of symbionts on unmeasured components of fitness while appropriately accounting for the uncertainty in these estimates.

The compiled dataset also provides insight into how data on host-symbiont interactions have been limited. We noted that particular taxa have dominated data collection. First, plant hosts made up the vast majority of the dataset, and preliminary screening of the search results indicated that significant additional effort would be required to generate an equivalent number of useable measurements of symbiotic effects on animal hosts (among the next 200 most relevant search results beyond those included in this analysis, only 9 studies focused on animal hosts).

The dataset did include insect and coral symbioses with microbes, however these interactions were less commonly manipulated experimentally, and measurements of performance tended to focus on fewer distinct life history processes. Even within the plant hosts, measurements came predominantly from relatively few well-known symbioses. Finally, the majority of measurements were taken in lab or greenhouse settings, meaning that measurements were most common for life stages amenable to experimentation. Measurements of growth for juvenile stages, typically recorded over the span of a few months, were the most common. While we did not find that measurements in field conditions differed from controlled settings (Fig S4 – S8), we expect that measurements later in the life cycle will be important for understanding how symbioses influence biological processes including senescence, lifetime reproductive output, and population age/stage structure. Expanding research that experimentally manipulates and measures symbioses in field environments will be particularly important to understand the responses of hosts and symbioses to global change (Kivlin et al. 2013; Rudgers et al. 2020) because variation inherent to natural environments is a key part of understanding how microbial symbioses influence populations and communities.

Finally, our analysis is a step in answering the long-standing call issued by Kostitzin for improved estimates of the vital coefficients that can form the basis of future theoretical and empirical work. Our meta-regression model provides broadly applicable estimates of symbiotic effects across host taxa. In future theoretical explorations of host-symbiont interactions, we urge researchers to parameterize their models using these sorts of empirical estimates of interaction outcomes to set realistic priors. We found that the magnitude of fitness effects of symbiosis is often consistent across different fitness components and life history stages, but that focused investment in key vital rates can be common for a given taxa. Key steps in the future will be investigating how variation in fitness effects interact with other mechanisms known to be important for the ecology and evolution of host-symbiont interactions, including transmission mode (Fine 1975; Genkai-Kato and Yamamura 1999; Afkhami and Rudgers 2008), inter-patch dispersal (Mihaljevic 2012; Mestre et al. 2020), and the potential for local adaptation in host and symbiont (De Mazancourt et al. 2005; Johnson et al. 2010). Our analysis of the diverse symbionts of grasses, which differ in their transmission mode (among other differences), showed showed different patterns of benefit across host life history, but this comparison across broad symbiont types conceals what is likely to be interesting variation and complexity of transmission. For example, does transmission depend on environmental context, and how commonly do mixed modes of transmission (both horizontal and vertical) occur? Empirical data on transmission rates are limited. Additionally, studies of multi-partner symbioses, beyond the pairwise interactions we evaluated here, are another important dimension of context-dependence to explore, as the presence of multiple symbionts (Afkhami et al. 2021; Palmer et al. 2010), or nested symbioses (Márquez et al. 2007; Batstone 2022), have been shown to have dramatic effects on interaction outcomes, and in a host community context, differential fitness benefits for hosts associating with different symbionts has the potential to mediate community-level outcomes (Kandlikar et al. 2019; Kandlikar et al. 2021). While there has been a push to develop mechanistic understanding of host-microbe symbioses (Johnson et al. 2013) by examining potential biotic and abiotic drivers of context dependence (Hoeksema and Bruna 2015) and how host functional traits may explain interaction outcomes (Gibert et al. 2019), we believe that embedding knowledge of symbiotic effects into demographic life history theory is the way forward for understanding how symbionts influence host populations.

## Summary

Previous theoretical work has suggested that conflicting effects of symbiosis on host vital rates is an important mechanism by which mutualisms have maintained evolutionary stability. Through a meta-analysis of published measurements of symbiotic effects on hosts, we found that symbiotic effects did not commonly occur in conflicting directions. While these effects were consistent in direction, we also documented strong outcomes of symbiosis for particular vital rates and stages that differed among host taxa and symbiont type. This empirical evidence suggests a more complicated understanding of mechanisms supporting mutualistic symbioses; symbionts alter hosts’ available resources and the magnitude of interaction effects is shaped by host and symbiont life history. Microbial symbiosis have great potential to influence population and community dynamics of hosts, but the meta-analytic dataset also show that there is clear need to explore how symbioses impact hosts beyond well-studied model systems and on older life stages. Synthesis of symbiont effects across diverse taxa revealed surprising consistency in outcomes, and variation across host life history serves as a basis for developing predictive understanding of host-symbiont dynamics.

## Acknowledgements

We thank the members of the Searcy and Afkhami labs who provided feedback on earlier versions of this research. This work was supported by funding from the National Science foundation (NSF-DEB 1922521, 2030060, and 2505581 to Michelle Afkhami and Chris Searcy, and an NSF Postdoctoral Fellowship 2410282 to Joshua Fowler).

## Data Accessibility Statement

Data for this publication is attached as supplemental material for peer-review and will be made available upon publication in a publicly accessible repository. Code to reproduce all analyses have been made available as an anonymized Zenodo repo (https://doi.org/10.5281/zenodo.21810630) for peer-review, and will be permanently archived in a publicly accessible repository and through github upon publication.

## Author Contributions

JCF contributed to research conception, data collection, data analysis, and led manuscript drafting. GP contributed to research conception, data collection, data analysis, and manuscript drafting. EH, VWL, ZM, and ALR contributed to data collection and manuscript drafting. CS and MA contributed to research conception, data analysis, and manuscript drafting.

## Conflict of Interest Statement

None Declared

## Notes

### Competing Interest Statement

The authors have declared no competing interest.

## References

1. Afkhami, Michelle E., Maren L. Friesen, and John R. Stinchcombe. 2021. “Multiple Mutualism Effects Generate Synergistic Selection and Strengthen Fitness Alignment in the Interaction between Legumes, Rhizobia and Mycorrhizal Fungi.” Ecology Letters 24 (9): 1824–34. 10.1111/ele.13814.

2. Afkhami, Michelle E., Patrick J. McIntyre, and Sharon Y. Strauss. 2014. “Mutualist-Mediated Effects on Species’ Range Limits across Large Geographic Scales.” Ecology Letters 17 (10): 1265–73. 10.1111/ele.12332.

3. Afkhami, Michelle E, and Jennifer A Rudgers. 2008. “Symbiosis Lost: Imperfect Vertical Transmission of Fungal Endophytes in Grasses.” The American Naturalist 172 (3): 405– 16. 10.1086/589893.

4. Alvarenga, Danillo O, and Kathrin Rousk. 2022. “Unraveling Host–Microbe Interactions and Ecosystem Functions in Moss–Bacteria Symbioses.” Journal of Experimental Botany 73 (13): 4473–86. 10.1093/jxb/erac091.

5. Angilletta Jr., Michael J., Christopher E. Oufiero, and Adam D. Leaché. 2006. “Direct and Indirect Effects of Environmental Temperature on the Evolution of Reproductive Strategies: An Information Theoretic Approach.” The American Naturalist 168 (4): E123–35. 10.1086/507880.

6. Armas, Cristina, Ramón Ordiales, and Francisco I. Pugnaire. 2004. “Measuring Plant Interactions: A New Comparative Index.” Ecology 85 (10): 2682–86. 10.1890/03-0650.

7. Batstone, Rebecca T. 2022. “Genomes within Genomes: Nested Symbiosis and Its Implications for Plant Evolution.” New Phytologist 234 (1): 28–34. 10.1111/nph.17847.

8. Bolan, N. S. 1991. “A Critical Review on the Role of Mycorrhizal Fungi in the Uptake of Phosphorus by Plants.” Plant and Soil 134 (2): 189–207. 10.1007/BF00012037.

9. Brucker, Robert M., and Seth R. Bordenstein. 2013. “The Hologenomic Basis of Speciation: Gut Bacteria Cause Hybrid Lethality in the Genus Nasonia.” Science 341 (6146): 667–69. 10.1126/science.1240659.

10. Bruner-Montero, Gaspar, and Francis M. Jiggins. 2023. “*Wolbachia* Protects *Drosophila Melanogaster* against Two Naturally Occurring and Virulent Viral Pathogens.” Scientific Reports 13 (1): 8518. 10.1038/s41598-023-35726-z.

11. Bürkner, Paul-Christian. 2017. “Brms: An R Package for Bayesian Multilevel Models Using Stan.” Journal of Statistical Software 80 (August): 1–28. 10.18637/jss.v080.i01.

12. Caswell, Hal. 2001. Matrix Population Models. Vol. 1. Sinauer Sunderland, MA, USA.

13. Chamberlain, Scott A., Judith L. Bronstein, and Jennifer A. Rudgers. 2014. “How Context Dependent Are Species Interactions?” Ecology Letters 17 (7): 881–90. 10.1111/ele.12279.

14. Chevalier, Mathieu, and Jonas Knape. 2020. “The Cost of Complexity in Forecasts of Population Abundances Is Reduced but Not Eliminated by Borrowing Information across Space Using a Hierarchical Approach.” Oikos 129 (2): 249–60. 10.1111/oik.06401.

15. Crouse, Deborah T., Larry B. Crowder, and Hal Caswell. 1987. “A Stage-Based Population Model for Loggerhead Sea Turtles and Implications for Conservation.” Ecology 68 (5): 1412–23. 10.2307/1939225.

16. De Mazancourt, Claire, Michel Loreau, and Ulf Dieckmann. 2005. “Understanding Mutualism When There Is Adaptation to the Partner.” Journal of Ecology 93 (2): 305–14. 10.1111/j.0022-0477.2004.00952.x.

17. Decunta, Facundo A., Luis I. Pérez, Dariusz P. Malinowski, Marco A. Molina-Montenegro, and Pedro E. Gundel. 2021. “A Systematic Review on the Effects of *Epichloë* Fungal Endophytes on Drought Tolerance in Cool-Season Grasses.” Frontiers in Plant Science 12: 380. 10.3389/fpls.2021.644731.

18. Donald, Marion L., Teresa F. Bohner, Kory M. Kolis, R. Alan Shadow, Jennifer A. Rudgers, and Tom E. X. Miller. 2021. “Context-Dependent Variability in the Population Prevalence and Individual Fitness Effects of Plant–Fungal Symbiosis.” Journal of Ecology 109 (2): 847–59. 10.1111/1365-2745.13510.

19. Egger, Matthias, George Davey Smith, Martin Schneider, and Christoph Minder. 1997. “Bias in Meta-Analysis Detected by a Simple, Graphical Test.” Paper. BMJ 315 (7109): 629–34. 10.1136/bmj.315.7109.629.

20. Ferriere, Régis, Judith L. Bronstein, Sergio Rinaldi, Richard Law, and Mathias Gauduchon. 2002. “Cheating and the Evolutionary Stability of Mutualisms.” Proceedings of the Royal Society of London. Series B: Biological Sciences 269 (1493): 773–80. 10.1098/rspb.2001.1900.

21. Fine, Paul E. M. 1975. “Vectors and Vertical Transmission: An Epidemiologic Perspective.” Annals of the New York Academy of Sciences 266 (1): 173–94. 10.1111/j.1749-6632.1975.tb35099.x.

22. Fowler, Joshua C., Marion L. Donald, Judith L. Bronstein, and Tom E. X. Miller. 2023. “The Geographic Footprint of Mutualism: How Mutualists Influence Species’ Range Limits.” Ecological Monographs 93 (1): e1558. 10.1002/ecm.1558.

23. Fowler, Joshua C., Shaun Ziegler, Kenneth D. Whitney, Jennifer A. Rudgers, and Tom E. X. Miller. 2024. “Microbial Symbionts Buffer Hosts from the Demographic Costs of Environmental Stochasticity.” Ecology Letters 27 (5): e14438. 10.1111/ele.14438.

24. Fukui, Shin. 2014. “Evolution of Symbiosis with Resource Allocation from Fecundity to Survival.” Die Naturwissenschaften 101 (5): 437–46. 10.1007/s00114-014-1175-1.

25. Gelman, Andrew, Xiao-Li Meng, and Hal Stern. 1996. “Posterior Predictive Assessment of Model Fitness Via Realized Discrepancies.” Statistica Sinica 6 (4): 733–60.

26. Gelman, Andrew, and Donald B. Rubin. 1992. “Inference from Iterative Simulation Using Multiple Sequences.” Statistical Science 7 (4): 457–72. 10.1214/ss/1177011136.

27. Gelman, Andrew, Daniel Simpson, and Michael Betancourt. 2017. “The Prior Can Often Only Be Understood in the Context of the Likelihood.” Entropy 19 (10): 10. 10.3390/e19100555.

28. Genkai-Kato, Motomi, and Norio Yamamura. 1999. “Evolution of Mutualistic Symbiosis without Vertical Transmission.” Theoretical Population Biology 55 (3): 309–23. 10.1006/tpbi.1998.1407.

29. Gibert, Anais, Wade Tozer, and Mark Westoby. 2019. “Plant Performance Response to Eight Different Types of Symbiosis.” *New Phytologist*, ahead of print, January. 10.1111/nph.15392.

30. Glick, Bernard R. 2012. “Plant Growth-Promoting Bacteria: Mechanisms and Applications.” Scientifica 2012 (1): 963401. 10.6064/2012/963401.

31. Guerrero, Ricardo, Lynn Margulis, and Mercedes Berlanga. 2013. “Symbiogenesis: The Holobiont as a Unit of Evolution.” Int Microbiol 16 (3): 133–43.

32. Gundel, P. E., P. H. Maseda, M. M. Vila-Aiub, C. M. Ghersa, and R. Benech-Arnold. 2006. “Effects of *Neotyphodium* Fungi on *Lolium multiflorum* Seed Germination in Relation to Water Availability.” Annals of Botany 97 (4): 571–77. 10.1093/aob/mcl004.

33. Hoang, Kim L., Heidi Choi, Nicole M. Gerardo, and Levi T. Morran. 2022. “Coevolution’s Conflicting Role in the Establishment of Beneficial Associations.” Evolution 76 (5): 1073–81. 10.1111/evo.14472.

34. Hoeksema, Jason D., and Emilio M. Bruna. 2015. “Context-Dependent Outcomes of Mutualistic Interactions.” Mutualism 10: 181–202.

35. Johnson, Nancy C., Caroline Angelard, Ian R. Sanders, and E. Toby Kiers. 2013. “Predicting Community and Ecosystem Outcomes of Mycorrhizal Responses to Global Change.” Ecology Letters 16 (s1): 140–53. 10.1111/ele.12085.

36. Johnson, Nancy Collins, Gail W. T. Wilson, Matthew A. Bowker, Jacqueline A. Wilson, and R. Michael Miller. 2010. “Resource Limitation Is a Driver of Local Adaptation in Mycorrhizal Symbioses.” Proceedings of the National Academy of Sciences 107 (5): 2093–98. 10.1073/pnas.0906710107.

37. Kandlikar, Gaurav S., Christopher A. Johnson, Xinyi Yan, Nathan J. B. Kraft, and Jonathan M. Levine. 2019. “Winning and Losing with Microbes: How Microbially Mediated Fitness Differences Influence Plant Diversity.” Ecology Letters 22 (8): 1178–91. 10.1111/ele.13280.

38. Kandlikar (□□□□□□□□□□□□), Gaurav S., Xinyi Yan (严心怡), Jonathan M. Levine, and Nathan J. B. Kraft. 2021. “Soil Microbes Generate Stronger Fitness Differences than Stabilization among California Annual Plants.” The American Naturalist 197 (1): E30–39. 10.1086/711662.

39. Kivlin, Stephanie N., Sarah M. Emery, and Jennifer A. Rudgers. 2013. “Fungal Symbionts Alter Plant Responses to Global Change.” American Journal of Botany 100 (7): 1445–57. 10.3732/ajb.1200558.

40. Kostitzin, V. A. (1934) 1978. “Symbiose, Parasitisme et Évolution: Étude Mathématique.” In The Golden Age of Theoretical Ecology, 1923-1940: A Collection of Works by V. Volterra, V.A. Kostitzin, A.J. Lotka, and A.N. Kolmogoroff, 22nd ed., edited by Francesco Scudo and Simon Levin. Lecture Notes in Biomathematics. 22nd ed. Springer-Verlag. https://cir.nii.ac.jp/crid/1130282271724708736.

41. Luo, Shan, Richard P. Phillips, Insu Jo, et al. 2023. “Higher Productivity in Forests with Mixed Mycorrhizal Strategies.” Nature Communications 14 (1): 1377. 10.1038/s41467-023-36888-0.

42. Macaskill, Petra, Stephen D. Walter, and Les Irwig. 2001. “A Comparison of Methods to Detect Publication Bias in Meta-Analysis.” Statistics in Medicine 20 (4): 641–54. 10.1002/sim.698.

43. Márquez, Luis M., Regina S. Redman, Russell J. Rodriguez, and Marilyn J. Roossinck. 2007. “A Virus in a Fungus in a Plant: Three-Way Symbiosis Required for Thermal Tolerance.” Science 315 (5811): 513–15. 10.1126/science.1136237.

44. Martinez, Fernando D. 2014. “The Human Microbiome. Early Life Determinant of Health Outcomes.” Annals of the American Thoracic Society 11 (Supplement 1): S7–12. 10.1513/AnnalsATS.201306-186MG.

45. Martinez, Julien, Suzan Ok, Sophie Smith, Kiana Snoeck, Jon P. Day, and Francis M. Jiggins. 2015. “Should Symbionts Be Nice or Selfish? Antiviral Effects of Wolbachia Are Costly but Reproductive Parasitism Is Not.” PLOS Pathogens 11 (7): e1005021. 10.1371/journal.ppat.1005021.

46. Mathew, Suni Anie, Marjo Helander, Kari Saikkonen, et al. 2023. “*Epichloë* Endophytes Shape the Foliar Endophytic Fungal Microbiome and Alter the Auxin and Salicylic Acid Phytohormone Levels in Two Meadow Fescue Cultivars.” Journal of Fungi 9 (1): 90. 10.3390/jof9010090.

47. Mestre, Alexandre, Robert Poulin, and Joaquín Hortal. 2020. “A Niche Perspective on the Range Expansion of Symbionts.” Biological Reviews 95 (2): 491–516. 10.1111/brv.12574.

48. Michonneau, François, Joseph W. Brown, and David J. Winter. 2016. “Rotl: An R Package to Interact with the Open Tree of Life Data.” Methods in Ecology and Evolution 7 (12): 1476–81. 10.1111/2041-210X.12593.

49. Mihaljevic, Joseph R. 2012. “Linking Metacommunity Theory and Symbiont Evolutionary Ecology.” Trends in Ecology & Evolution 27 (6): 323–29. 10.1016/j.tree.2012.01.011.

50. Moreno, Santiago G., Alex J. Sutton, AE Ades, et al. 2009. “Assessment of Regression-Based Methods to Adjust for Publication Bias through a Comprehensive Simulation Study.” BMC Medical Research Methodology 9 (1): 2. 10.1186/1471-2288-9-2.

51. Nakagawa, Shinichi, Malgorzata Lagisz, Michael D. Jennions, et al. 2022. “Methods for Testing Publication Bias in Ecological and Evolutionary Meta-Analyses.” Methods in Ecology and Evolution 13 (1): 4–21. 10.1111/2041-210X.13724.

52. Olanrewaju, Oluwaseyi Samuel, Bernard R. Glick, and Olubukola Oluranti Babalola. 2017. “Mechanisms of Action of Plant Growth Promoting Bacteria.” World Journal of Microbiology and Biotechnology 33 (11): 197. 10.1007/s11274-017-2364-9.

53. Oliver, Kerry M., Jacob A. Russell, Nancy A. Moran, and Martha S. Hunter. 2003. “Facultative Bacterial Symbionts in Aphids Confer Resistance to Parasitic Wasps.” Proceedings of the National Academy of Sciences 100 (4): 1803–7. 10.1073/pnas.0335320100.

54. Palmer, Todd M., Daniel F. Doak, Maureen L. Stanton, et al. 2010. “Synergy of Multiple Partners, Including Freeloaders, Increases Host Fitness in a Multispecies Mutualism.” Biological Sciences. Proceedings of the National Academy of Sciences 107 (40): 17234– 39. 10.1073/pnas.1006872107.

55. Panaccione, Daniel G., Wesley T. Beaulieu, and Daniel Cook. 2014. “Bioactive Alkaloids in Vertically Transmitted Fungal Endophytes.” Functional Ecology 28 (2): 299–314. 10.1111/1365-2435.12076.

56. Petersen, Carola, Inga K Hamerich, Karen L Adair, et al. 2023. “Host and Microbiome Jointly Contribute to Environmental Adaptation.” The ISME Journal 17 (11): 1953–65. 10.1038/s41396-023-01507-9.

57. Pfister, Catherine A. 1998. “Patterns of Variance in Stage-Structured Populations: Evolutionary Predictions and Ecological Implications.” Proceedings of the National Academy of Sciences 95 (1): 213–18. 10.1073/pnas.95.1.213.

58. Pick, Joel L., Shinichi Nakagawa, and Daniel W. A. Noble. 2019. “Reproducible, Flexible and High-Throughput Data Extraction from Primary Literature: The metaDigitise r Package.” Methods in Ecology and Evolution 10 (3): 426–31. 10.1111/2041-210X.13118.

59. Plard, Floriane, Nigel G. Yoccoz, Christophe Bonenfant, François Klein, Claude Warnant, and Jean-Michel Gaillard. 2015. “Disentangling Direct and Growth-Mediated Influences on Early Survival: A Mechanistic Approach.” Journal of Animal Ecology 84 (5): 1363–72. 10.1111/1365-2656.12378.

60. Rodriguez, R. J., J. F. White Jr, A. E. Arnold, and R. S. Redman. 2009. “Fungal Endophytes: Diversity and Functional Roles.” New Phytologist 182 (2): 314–30. 10.1111/j.1469-8137.2009.02773.x.

61. Rosenberg, Eugene, Gil Sharon, Ilil Atad, and Ilana Zilber-Rosenberg. 2010. “The Evolution of Animals and Plants via Symbiosis with Microorganisms.” Environmental Microbiology Reports 2 (4): 500–506. 10.1111/j.1758-2229.2010.00177.x.

62. Rudgers, Jennifer A., Michelle E. Afkhami, Lukas Bell-Dereske, et al. 2020. “Climate Disruption of Plant-Microbe Interactions.” Annual Review of Ecology, Evolution, and Systematics 51 (1): null. 10.1146/annurev-ecolsys-011720-090819.

63. Rudgers, Jennifer A., Jennifer M. Koslow, and Keith Clay. 2004. “Endophytic Fungi Alter Relationships between Diversity and Ecosystem Properties.” Ecology Letters 7 (1): 42– 51. 10.1046/j.1461-0248.2003.00543.x.

64. Rudgers, Jennifer A., Tom E. X. Miller, Shaun M. Ziegler, and Kelly D. Craven. 2012. “There Are Many Ways to Be a Mutualist: Endophytic Fungus Reduces Plant Survival but Increases Population Growth.” Ecology 93 (3): 565–74. 10.1890/11-0689.1.

65. Ruiz-Lozano, J. M., and R. Azcón. 1995. “Hyphal Contribution to Water Uptake in Mycorrhizal Plants as Affected by the Fungal Species and Water Status.” Physiologia Plantarum 95 (3): 472–78. 10.1111/j.1399-3054.1995.tb00865.x.

66. Sæther, Bernt-Erik, Tim Coulson, Vidar Grøtan, et al. 2013. “How Life History Influences Population Dynamics in Fluctuating Environments.” The American Naturalist 182 (6): 743–59. 10.1086/673497.

67. Sarkar, Anujit, Ji Youn Yoo, Samia Valeria Ozorio Dutra, Katherine H. Morgan, and Maureen Groer. 2021. “The Association between Early-Life Gut Microbiota and Long-Term Health and Diseases.” Journal of Clinical Medicine 10 (3): 459. 10.3390/jcm10030459.

68. Schaub, M., and M. Kéry. 2012. “Combining Information in Hierarchical Models Improves Inferences in Population Ecology and Demographic Population Analyses.” Animal Conservation 15 (2): 125–26. 73931168. 10.1111/j.1469-1795.2012.00531.x.

69. Schwarzer, Guido, James R. Carpenter, and Gerta Rücker. 2015. “Small-Study Effects in Meta-Analysis.” In Meta-Analysis with R, edited by Guido Schwarzer, James R. Carpenter, and Gerta Rücker. Springer International Publishing. 10.1007/978-3-319-21416-0_5.

70. Siddaway, Andy P., Alex M. Wood, and Larry V. Hedges. 2019. “How to Do a Systematic Review: A Best Practice Guide for Conducting and Reporting Narrative Reviews, Meta-Analyses, and Meta-Syntheses.” Annual Review of Psychology 70 (Volume 70, 2019): 747–70. 10.1146/annurev-psych-010418-102803.

71. Simon, Jean-Christophe, Julian R. Marchesi, Christophe Mougel, and Marc-André Selosse. 2019. “Host-Microbiota Interactions: From Holobiont Theory to Analysis.” Microbiome 7 (1): 5. 10.1186/s40168-019-0619-4.

72. Stearns, S. C. 1989. “Trade-Offs in Life-History Evolution.” Functional Ecology 3 (3): 259–68. 10.2307/2389364.

73. Sterne, Jonathan A. C., Matthias Egger, and George Davey Smith. 2001. “Investigating and Dealing with Publication and Other Biases in Meta-Analysis.” Education and Debate. BMJ 323 (7304): 101–5. 10.1136/bmj.323.7304.101.

74. Stott, Iain, Roberto Salguero-Gómez, Owen R. Jones, et al. 2024. “Life Histories Are Not Just Fast or Slow.” Trends in Ecology & Evolution 39 (9): 830–40. 10.1016/j.tree.2024.06.001.

75. Subedi, Suresh C., Preston Allen, Rosario Vidales, Leonel Sternberg, Michael Ross, and Michelle E. Afkhami. 2022. “Salinity Legacy: Foliar Microbiome’s History Affects Mutualist-Conferred Salinity Tolerance.” Ecology 103 (6): e3679. 10.1002/ecy.3679.

76. Thompson, Simon G., and Stephen J. Sharp. 1999. “Explaining Heterogeneity in Meta-Analysis: A Comparison of Methods.” Statistics in Medicine 18 (20): 2693–708. 10.1002/(SICI)1097-0258(19991030)18:20%253C2693::AID-SIM235%253E3.0.CO;2-V.

77. Travis, J. M. J., R. W. Brooker, E. J. Clark, and C. Dytham. 2006. “The Distribution of Positive and Negative Species Interactions across Environmental Gradients on a Dual-Lattice Model.” Journal of Theoretical Biology 241 (4): 896–902. 10.1016/j.jtbi.2006.01.025.

78. Vehtari, Aki, Andrew Gelman, Daniel Simpson, Bob Carpenter, and Paul-Christian Bürkner. 2021. “Rank-Normalization, Folding, and Localization: An Improved R^ for Assessing Convergence of MCMC (with Discussion).” Bayesian Analysis 16 (2): 667–718. 10.1214/20-BA1221.

79. Waqas, Muhammad, Abdul Latif Khan, Muhammad Kamran, et al. 2012. “Endophytic Fungi Produce Gibberellins and Indoleacetic Acid and Promotes Host-Plant Growth during Stress.” Molecules 17 (9): 10754–73. 10.3390/molecules170910754.

80. Yule, Kelsey M., Tom E. X. Miller, and Jennifer A. Rudgers. 2013. “Costs, Benefits, and Loss of Vertically Transmitted Symbionts Affect Host Population Dynamics.” Oikos 122 (10): no-no. 10.1111/j.1600-0706.2012.00229.x.

81. Zera, Anthony J., and Lawrence G. Harshman. 2001. “The Physiology of Life History Trade-Offs in Animals.” *Annual Review of Ecology*, Evolution, and Systematics 32 (Volume 32, 2001): 95–126. 10.1146/annurev.ecolsys.32.081501.114006.

82. Zhao, Zhenrui, Mingzhu Kou, Rui Zhong, Chao Xia, Michael J. Christensen, and Xingxu Zhang. 2021. “Transcriptome Analysis Revealed Plant Hormone Biosynthesis and Response Pathway Modification by *Epichloëgansuensis* in *Achnatheruminebrians* under Different Soil Moisture Availability.” Journal of Fungi 7 (8): 640. 10.3390/jof7080640.

