## Supplemental Methods, Figures, and Tables for "Do microbial effects on hosts vary across life stage and vital rate?"

**Supplemental Material**

Supplemental Methods

1. ***Meta-regression analysis of small study bias; Figures S1-S4***
2. ***Evaluating how symbiont effects differ across experimental contexts; Figures S3-S7***

Supplemental Figures S8-S15

Supplemental Tables S1-S2

**Supplemental Methods**

***Meta-regression analysis of small study bias***

To assess potential small study bias within the compiled dataset of host-symbiont interaction effects, we performed a regression analysis analogous to an Egger’s test (Egger et al. 1997; Schwarzer et al. 2015), but implemented in a Bayesian framework. An Egger’s test evaluates the relationship between scaled effect sizes and study precision, where an intercept different from zero suggests publication bias. The regression model describing an Egger’s test can be formulated as follows:

$$\begin{aligned} \frac{RII}{{SE}_{RII}}\sim Normal\left( \mu_{i},\sigma\right) \#\left( S.1 \right) \end{aligned}$$

$$\mu_{i}=\alpha+ \beta*(\frac{1}{{SE}_{RII}})$$

Where the relationship between the scaled effect size ($\frac{RII}{{SE}_{RII}}$) and the precision of this estimate ($\frac{1}{{SE}_{RII}})$is determined by an intercept $\alpha$, and slope $\beta$. Publication bias is suspected when the intercept term differs statistically from zero; positive intercept values suggest that studies with low precision overestimate true effects. In a classic meta-analysis, Egger’s test assumes that differences in effect size between low and high precision studies is solely due to publication bias and ignores other sources of heterogeneity. To accommodate known heterogeneity in our dataset, which included measurements across different vital rates, life stages, and host species, we constructed a mixed effects regression as follows:

$$\begin{aligned} \frac{RII_{i}}{{SE}_{RII}} \sim Normal\left( \mu_{i,s}, \sigma\right) \\ \mu_{i}=\alpha_{v}+ \beta_{v}*\left( \frac{1}{\mathrm{SE}_{\mathrm{RII}}} \right)+\tau\\ \alpha\sim Normal\left( 0,100 \right) \\ \beta\sim Normal\left( 0,100 \right) \\ \tau\sim Normal\left( 0,100 \right) \\ \sigma\sim half-Normal\left( 0,100 \right) \\ \\ \end{aligned}(S.2)$$

Where intercept ($\alpha_{v}$) and slope ($\beta_{v}$) parameters varied with vital rate and a study-level random effect, $\tau$, accounted for variation across studies that contain multiple measurements. To test for potential differences across different life stages, we repeated this regression analysis, allowing parameters to vary with life stage rather than vital rate. All parameters were given wide, Gaussian priors with mean = 0 and sd = 100. The variance parameter was given a positively defined, half-normal distribution prior with mode = 0, sd = 100.

This regression approach is similar to that presented in Nakagawa et al. 2021, which developed a Bayesian hierarchical meta-regression approach for bias tests in meta-analyses. Like Nakagawa et al. 2021, we account for heterogeneity among effect sizes in our analysis through the use of covariates and hierarchical (random) effects. A key difference is that this regression has as its dependent variable scaled effect sizes, as in the original formulation of Egger’s test, whereas Nakagawa et al. 2021 perform a weighted regression, such that estimates are weighted by the sampling variance. Previous work developing bias tests has shown the choice of weighted vs unweighted regression can influence the potential for false positives, and that different methods perform favorably on different effect size measurements (i.e., SMD, log(RR)) (Thompson and Sharp 1999; Macaskill et al. 2001; Sterne et al. 2001; Moreno et al. 2009). We chose an effect size measurement, RII, because of its other favorable statistical properties described in the main text (bounded between 1 and −1; and symmetrical around zero), but this effect size has not seen wide uptake outside of ecological research. Our goal in this bias analysis is primarily to identify potential biases, rather than adjust the central meta-analytic estimates to account for these biases. While these choices likely impact our interpretation of potential bias within the dataset, we expect that differences in statistical significance would be minimal, and that this analysis provides an interpretable and easily implemented regression that fits its purpose.

Overall, we did not find strong support for small study publication bias within the compiled dataset. However, the results of this analysis pointed to several small biases or errors in initial data collection. Initial plotting and model fitting identified several effect sizes with unusually high precision measurements, and we investigated these to ensure that they were not the result of data transcription errors and corrected three such results (in which sample sizes were recorded as a magnitude larger than was correct). Another set of measurements from one study had precision 100 times greater than other studies. Inclusion of these measurements in the bias test led to statistically significant negative intercept values, in the opposite direction classically interpreted as evidence of “small study bias”. While this standard deviation was correctly transcribed from a supplemental table, we believe it is likely these outlier precision values represent a typo or miscalculation on the part of the original authors, and so we removed these estimates from subsequent analyses. After this step, none of the intercept parameters across all the vital rates or life stages were statistically different from 0 (Figs. S1; S2), suggesting that there is not meaningful small study bias within our dataset of host-symbiont interactions. While the 95% credible interval in all cases overlapped zero, we do find that the estimated intercept parameters were marginally negative for some vital rate metrics, particularly for measurements of juvenile growth. Again, positive intercepts are classically interpreted as evidence of small publication bias (low precision studies tend to overestimate true effects), and this marginally negative intercept could suggest that some low precision studies of microbial effects on hosts underestimate symbiont effects for young life stages.

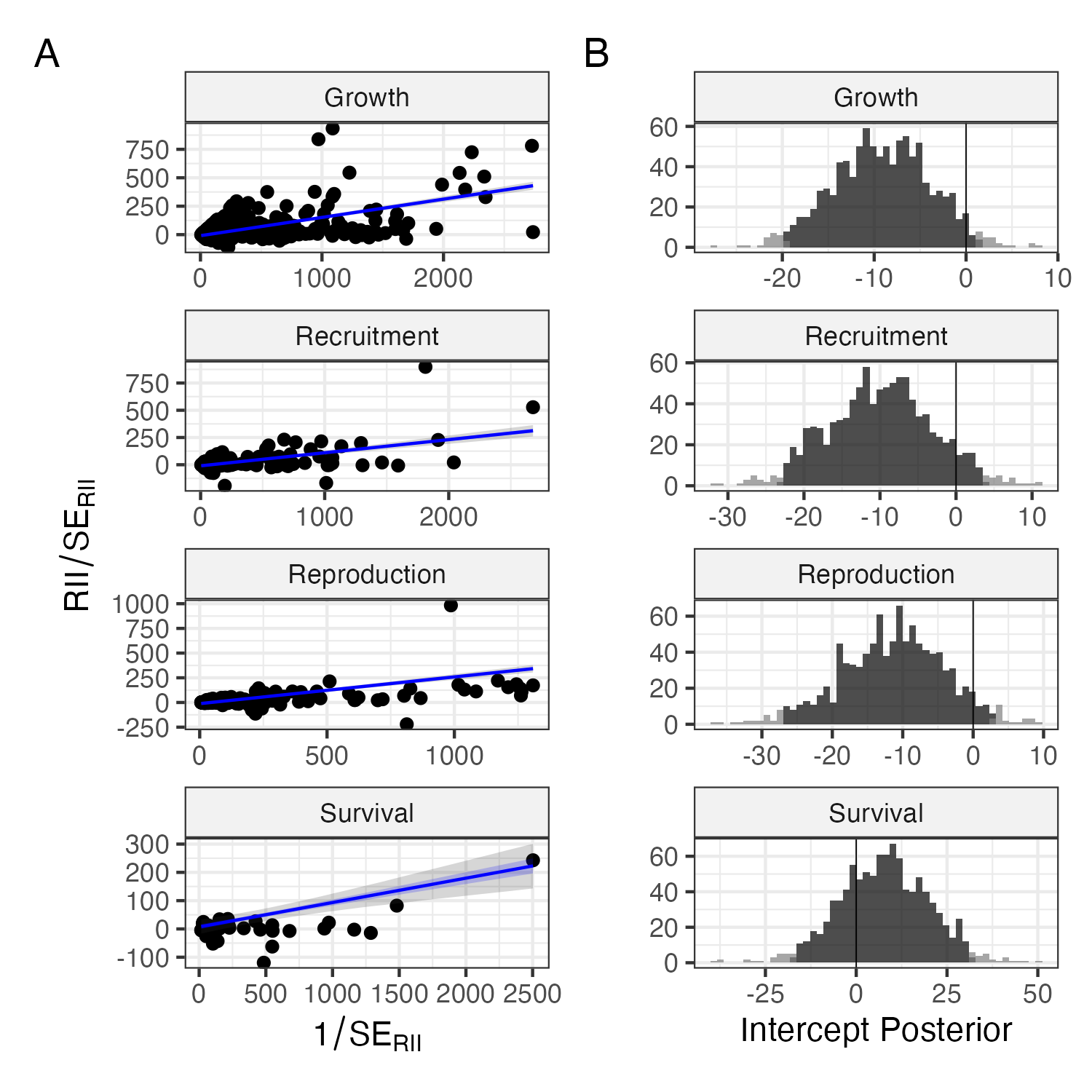

**Figure S1. Minimal publication bias within measured effect sizes across vital rates.** (A) Panels show the relationship between scaled effect sizes and measurement precision across vital rates. Points are values from each study; regression lines show the predicted mean along shaded bands depicting 50% and 95% CI. (B) Posterior distribution of regression intercepts for each vital rate. Dark shading within histograms depicts the 95% credible interval.

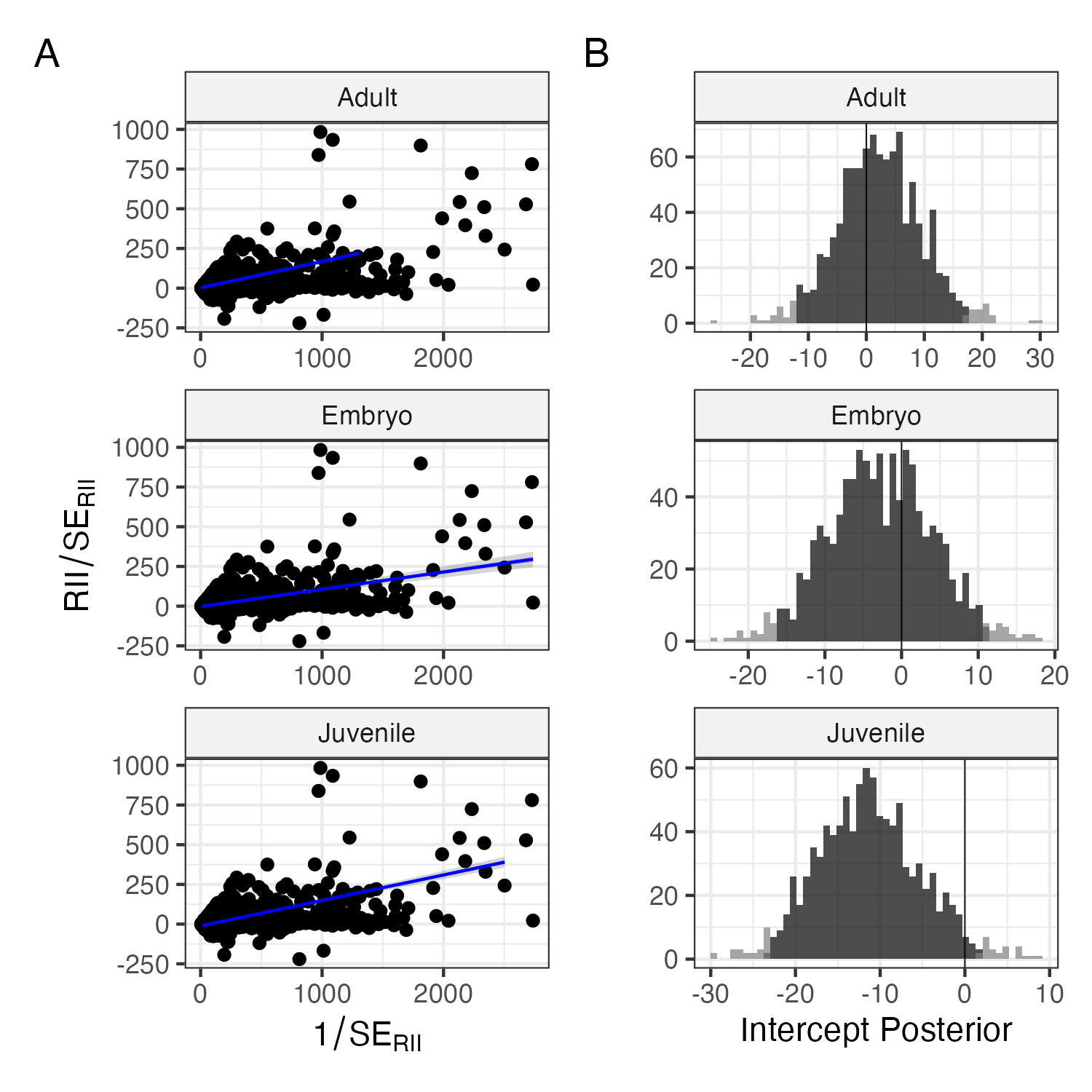

**Figure S2. Minimal publication bias within measured effect sizes across life stages.** (A) Panels show the relationship between scaled effect sizes and measurement precision across host life stages. Points are values from each study; regression lines show the predicted mean along shaded bands depicting 50% and 95% CI. (B) Posterior distribution of regression intercepts for each vital rate. Dark shading within histograms depicts the 95% credible interval.

***Evaluating how symbiont effects differ across experimental contexts***

To evaluate whether symbiotic effects differed across experimental settings, we again modeled RII with a set of meta-regression models, as described in the main text (Eqn. 2), where observed effect sizes, RII, assumed a Gaussian distribution with measurement error. We again performed separate analyses for differences across vital rates and differences across life stages. We incorporated interaction terms that allowed fixed parameters to vary across three experimental contexts (lab, greenhouse, or field environments), and also across experimental (involving the addition or removal of symbionts from hosts) or observational (comparison of naturally occurring symbiotic and aposymbiotic hosts) symbiont treatments. The linear predictor describing this model is given in Eqn. S.1:

$$\begin{aligned} \mu_{i} = \beta_{v,\left[ setting \right],[treatment]}+\tau_{f,v}+\gamma_{g|f,v} + \zeta_{s}+\eta_{e|s}\#\left( S.3 \right) \end{aligned}$$

where the intercept parameters ($\beta$) vary according to experimental setting and symbiont treatment for each vital rate descriptor (v). Nested random effects described variance associated with host taxonomy ($\tau,\gamma$) accounting for host family (*f*) and host genus (*g*) as well as study-level random effects ($\eta$ and $\zeta$) accounting for multiple separate measurements of effect size from each experiment (e) within each paper (s). Prior choices and model fitting were as described in the main text (Eqn. 2).

This analysis demonstrated, that after accounting for host taxonomy and study-level variance, there were no consistent differences between experimental contexts. Comparing between lab, greenhouse, and field settings, symbiont effects were similar within each vital rate (Fig. S3). In particular, we did not find that experimental manipulations were stronger or weaker than observational, or that they differed between field, lab and greenhouse contexts. Posterior estimates of the difference between experimental and observational measurements were consistently centered around zero (Fig. S5).

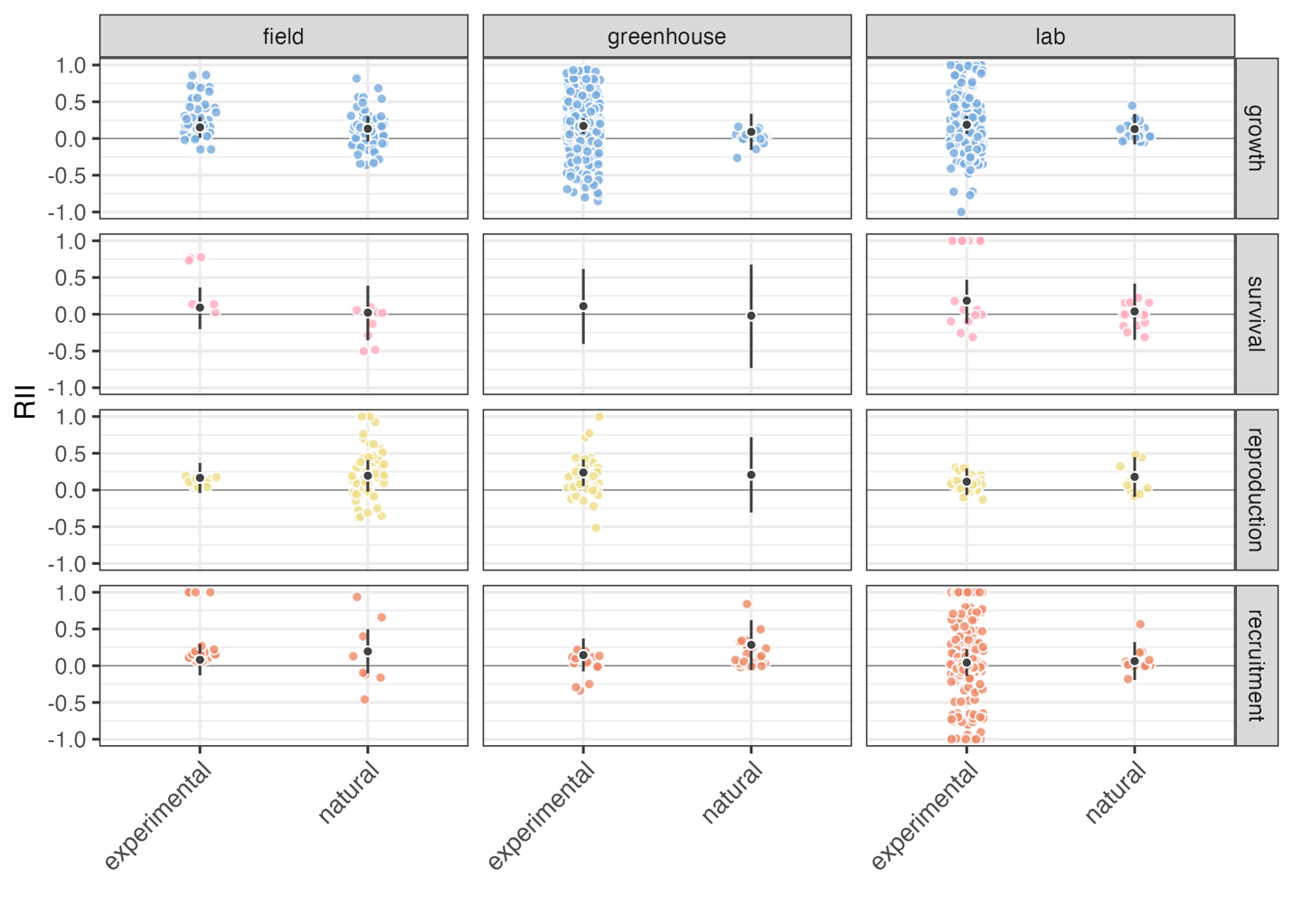

**Figure S4. No differences across experimental contexts in symbiont effects on host vital rates.** Meta-analytic means (black) along with 95% CI are shown for experimental and observational symbiont treatments for each vital rate type (growth, survival, reproduction, and recruitment) and experimental setting (field, greenhouse, and lab), along with observed RII values across studies (colored points).

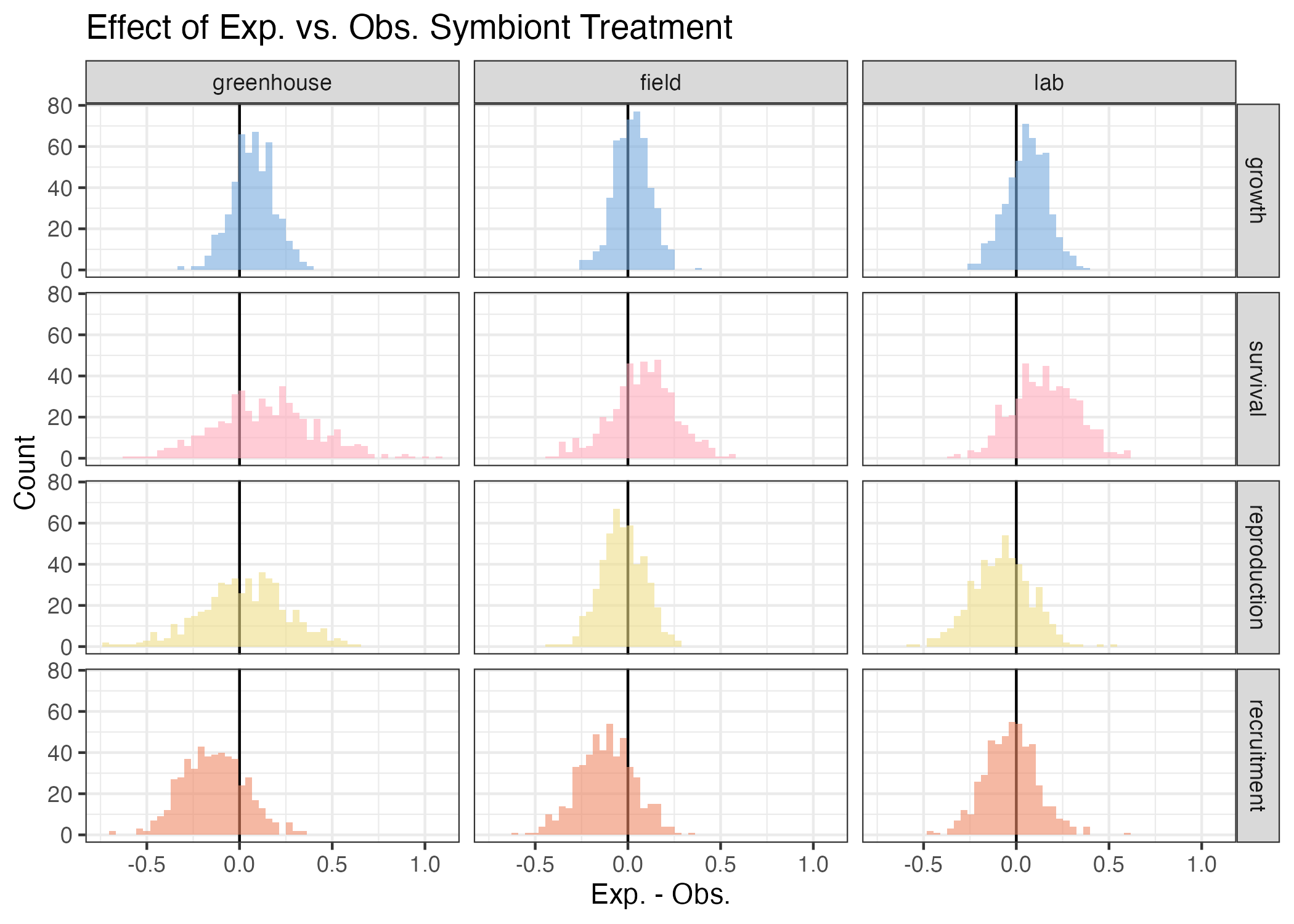

**Figure S5. Posterior distribution of the effect of experimental vs observational symbiont treatments across host vital rates. Each panel shows the posterior distribution from 500 posterior draws of the difference between experimental and observational studies for each vital rate (growth, survival, reproduction, and recruitment) and each experimental context (greenhouse, field, and lab).**

Similarly, we found that there were no consistent differences in the magnitude of symbiont effects across host life stages between experimental contexts. Comparing between lab, greenhouse, and field settings, symbiont effects were similar within each life stage (Fig. S6). We did not find that experimental manipulations were stronger or weaker than observational, or that they differed between field, lab, and greenhouse contexts. Posterior estimates of the difference between experimental and observational measurements were consistently centered around zero (Fig. S7).

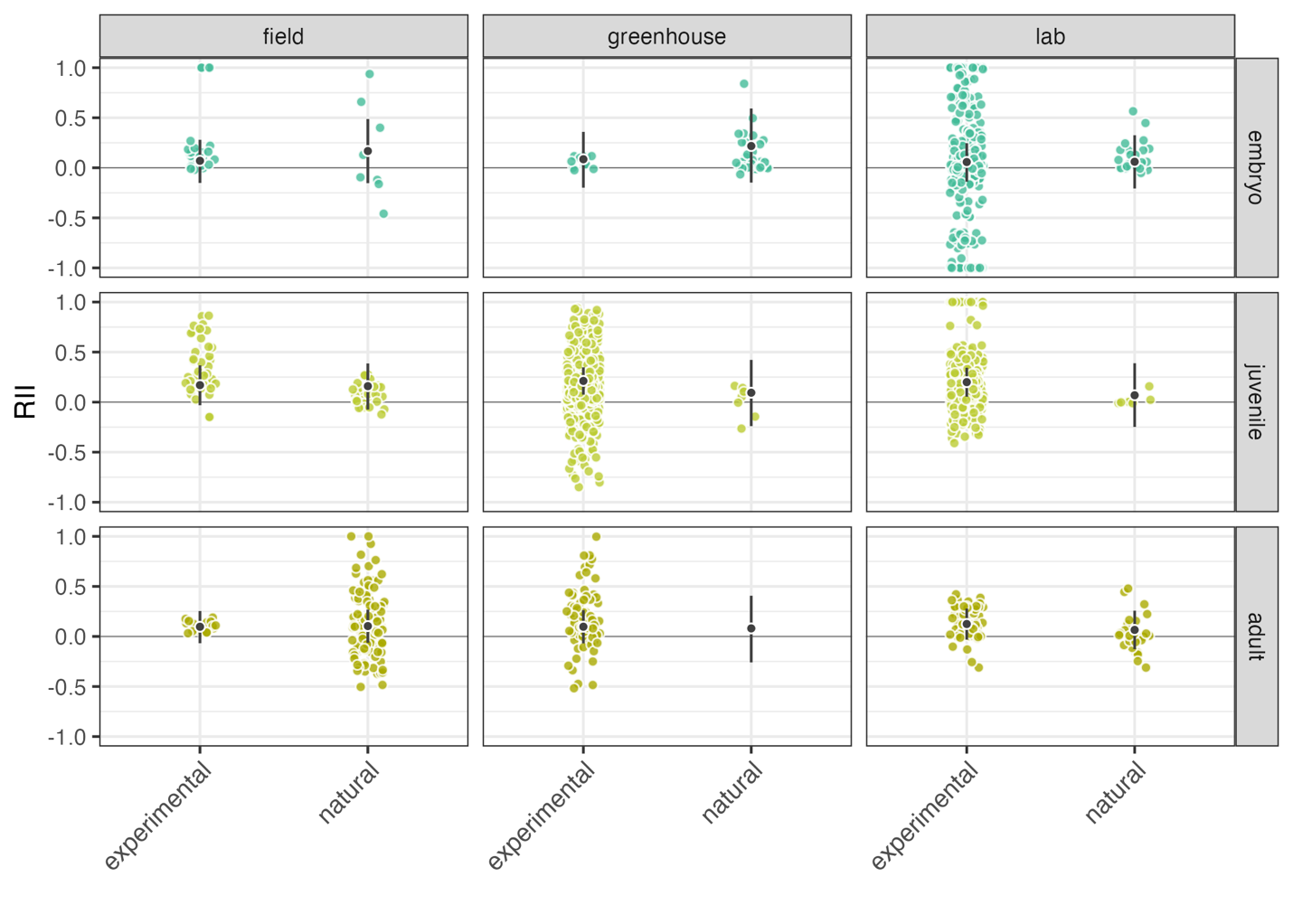

**Figure S6. No differences across experimental contexts in symbiont effects across host life stages.** Meta-analytic means (black) along with 95% CI are shown for experimental and observational symbiont treatments for each vital rate type (growth, survival, reproduction, and recruitment) and experimental setting (field, greenhouse, and lab), along with observed RII values across studies (colored points).

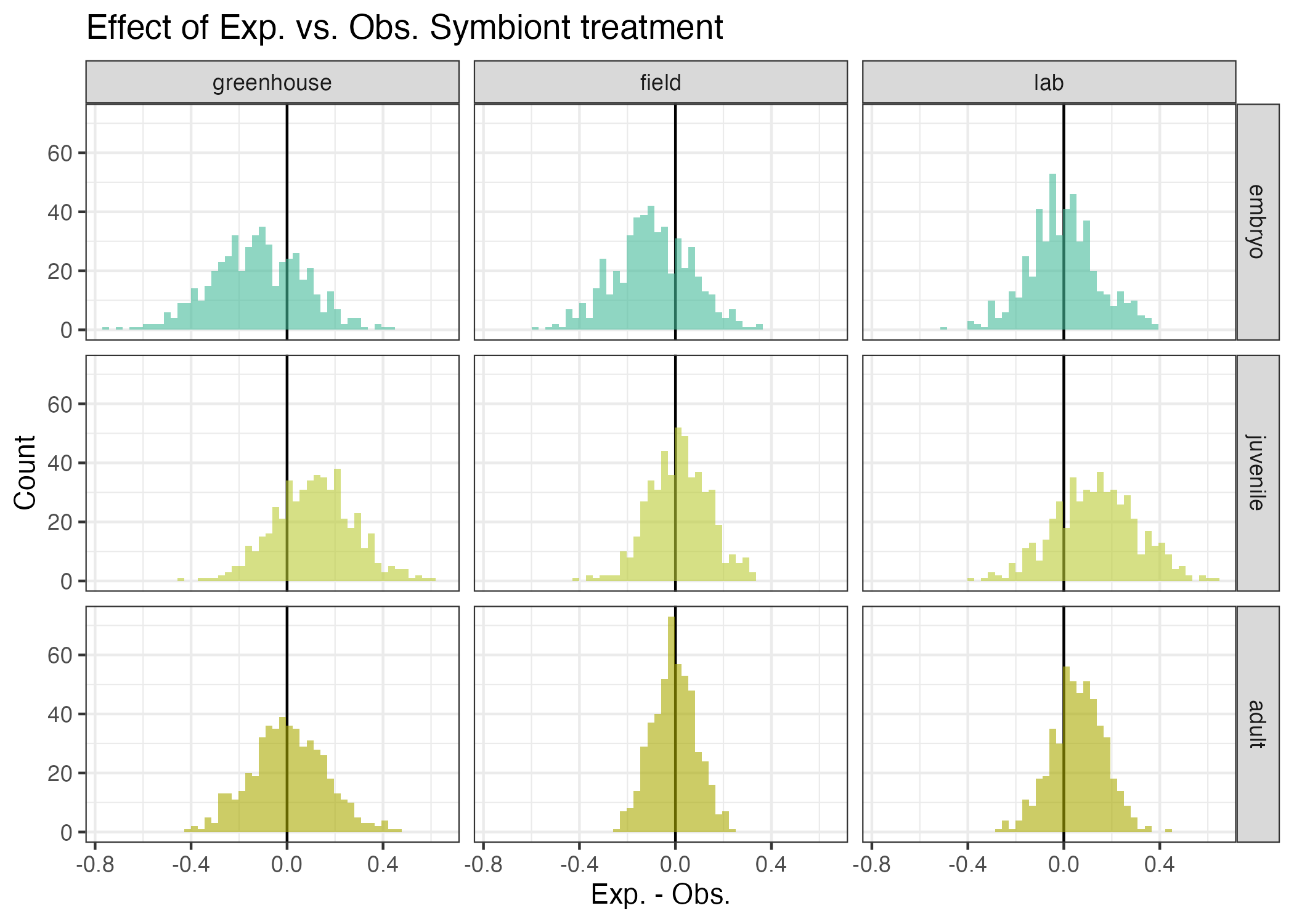

**Figure S7. Posterior distribution of the effect of experimental vs observational symbiont treatments across host life stages. Each panel shows the posterior distribution from 500 posterior draws of the difference between experimental and observational studies for each vital rate (growth, survival, reproduction, and recruitment) and each experimental context (greenhouse, field, and lab).**

**Supplemental Figures S8-S15**

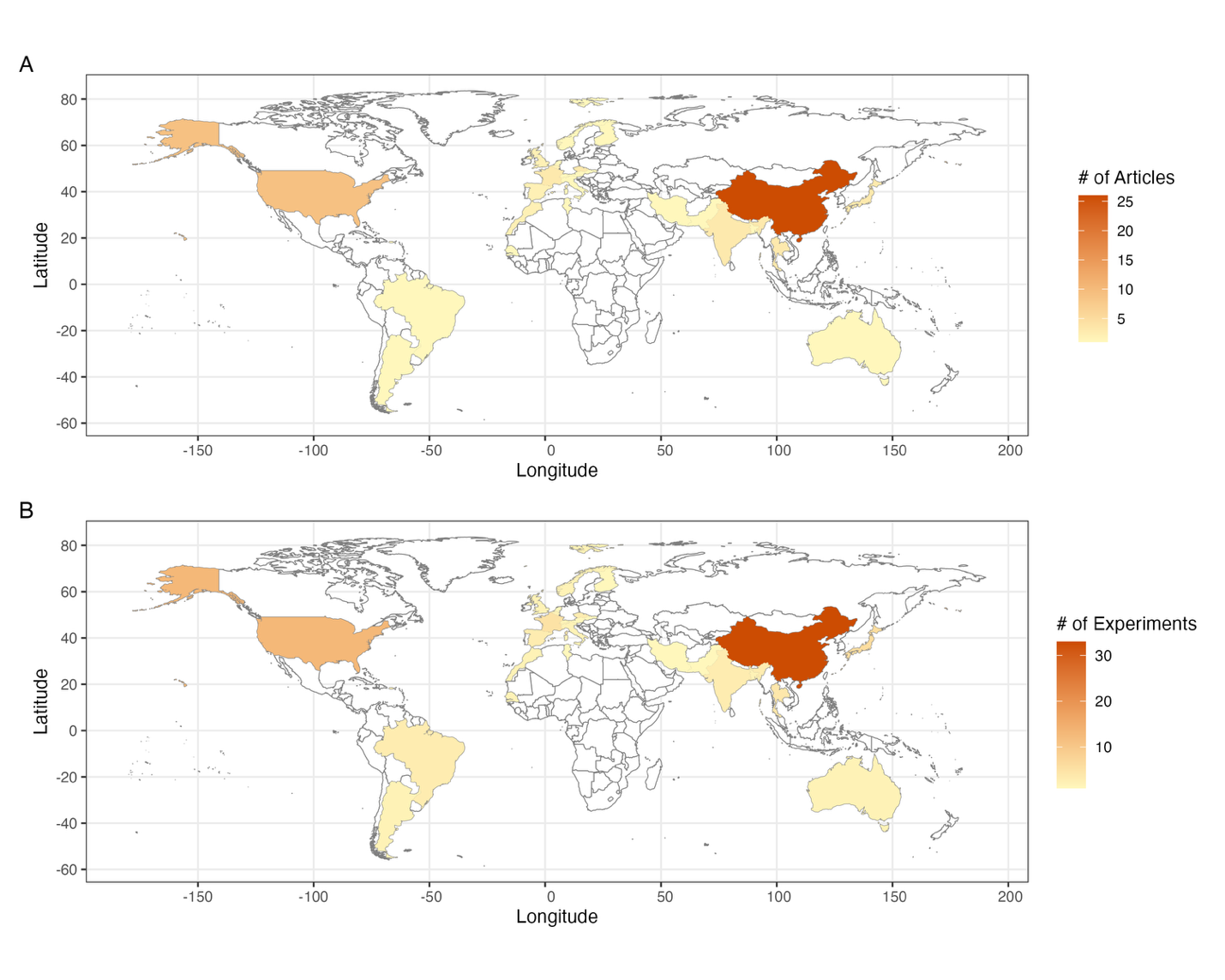

**Figure S8. Global distribution of data compiled during the literature search.** Across each country, shading indicates the number of unique (A) articles or (B) experiments within the meta-analytic dataset of measurements of host-symbiont interactions.

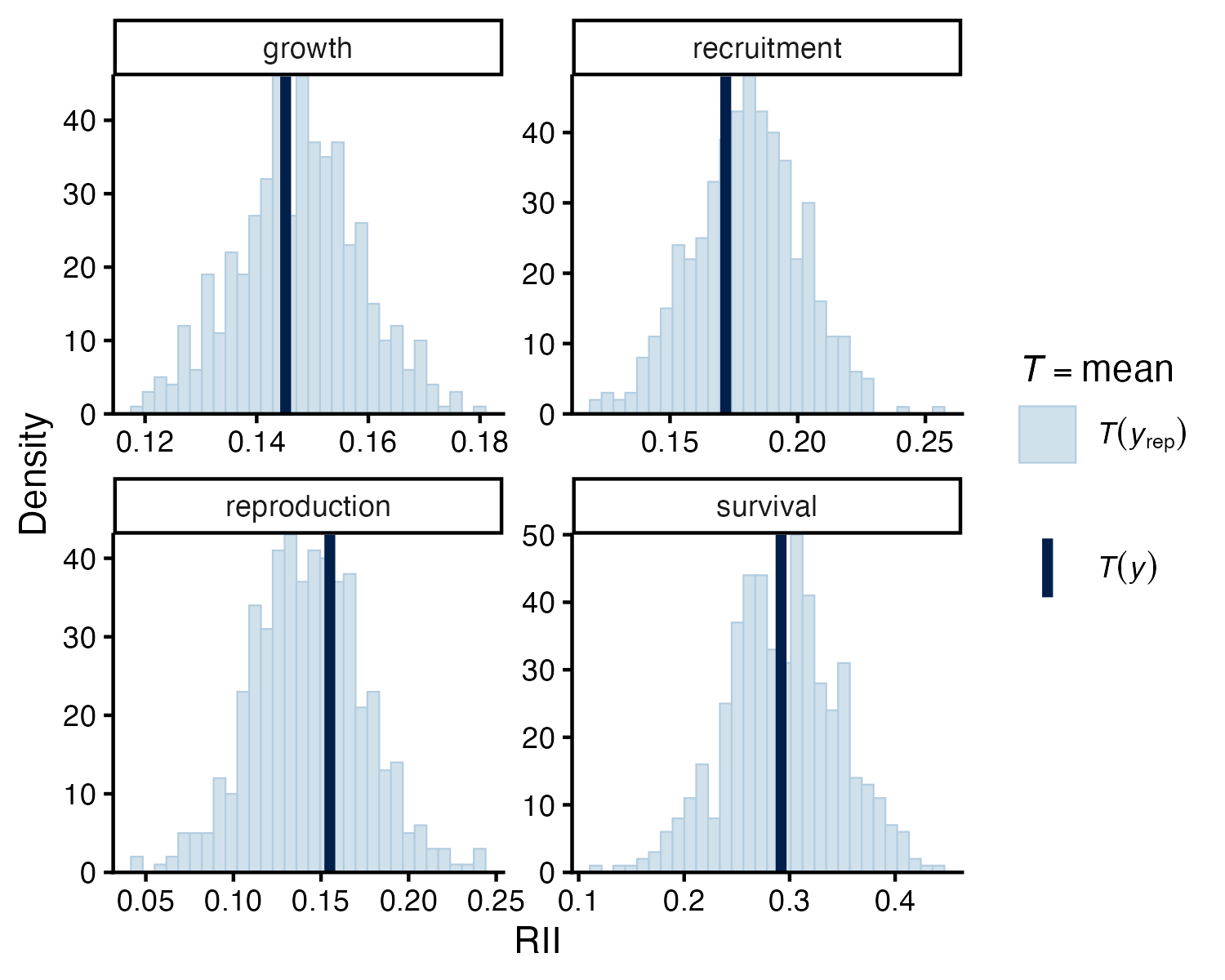

**Figure S9 Graphical posterior predictive check for meta-regression model of symbiotic effects on host vital rates.** Consistency between observed data and predicted data indicate models describe the data well. Dark blue bars show mean effect size (RII) of observed dataset within each vital rate category along with light blue histograms showing 500 predicted mean effect sizes from model posteriors.

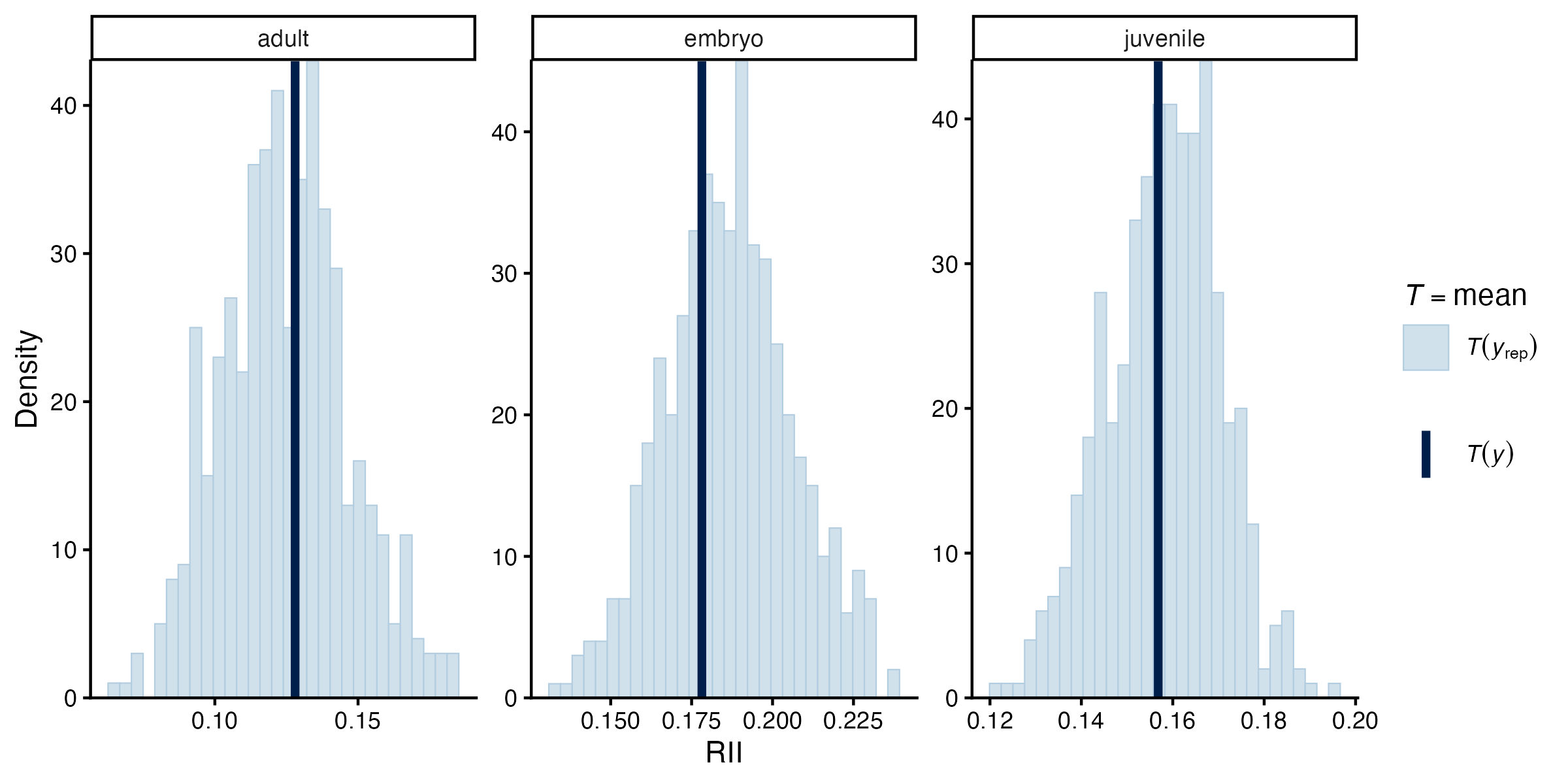

**Figure S10. Graphical posterior predictive check for meta-regression model of symbiotic effects across host life stages.** Consistency between observed data and predicted data indicate models describe the data well. Dark blue bars show mean effect size (RII) of observed dataset within each vital rate category along with light blue histograms showing 500 predicted mean effect sizes from model posteriors.

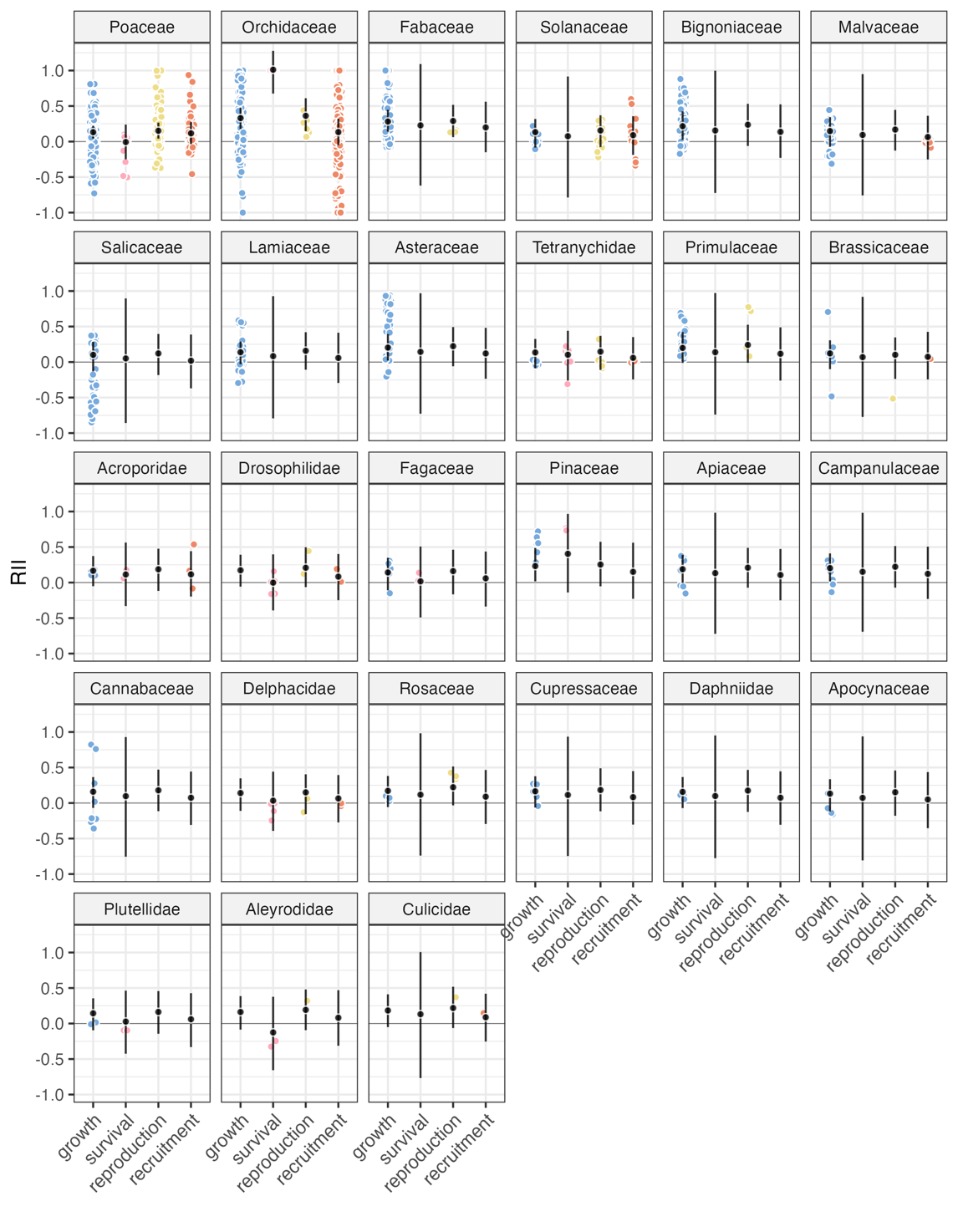

**Figure S11. Family specific meta-analytic mean estimates of microbial effects on host vital rates.** Meta-analytic means (black) along with 95% CI are shown for each vital rate type (growth, survival, reproduction, and recruitment), along with observed RII values across studies. Predictions averaging across taxonomic diversity are shown for each of 27 host families within the dataset. Panels for each family are presented in descending order of sample size.

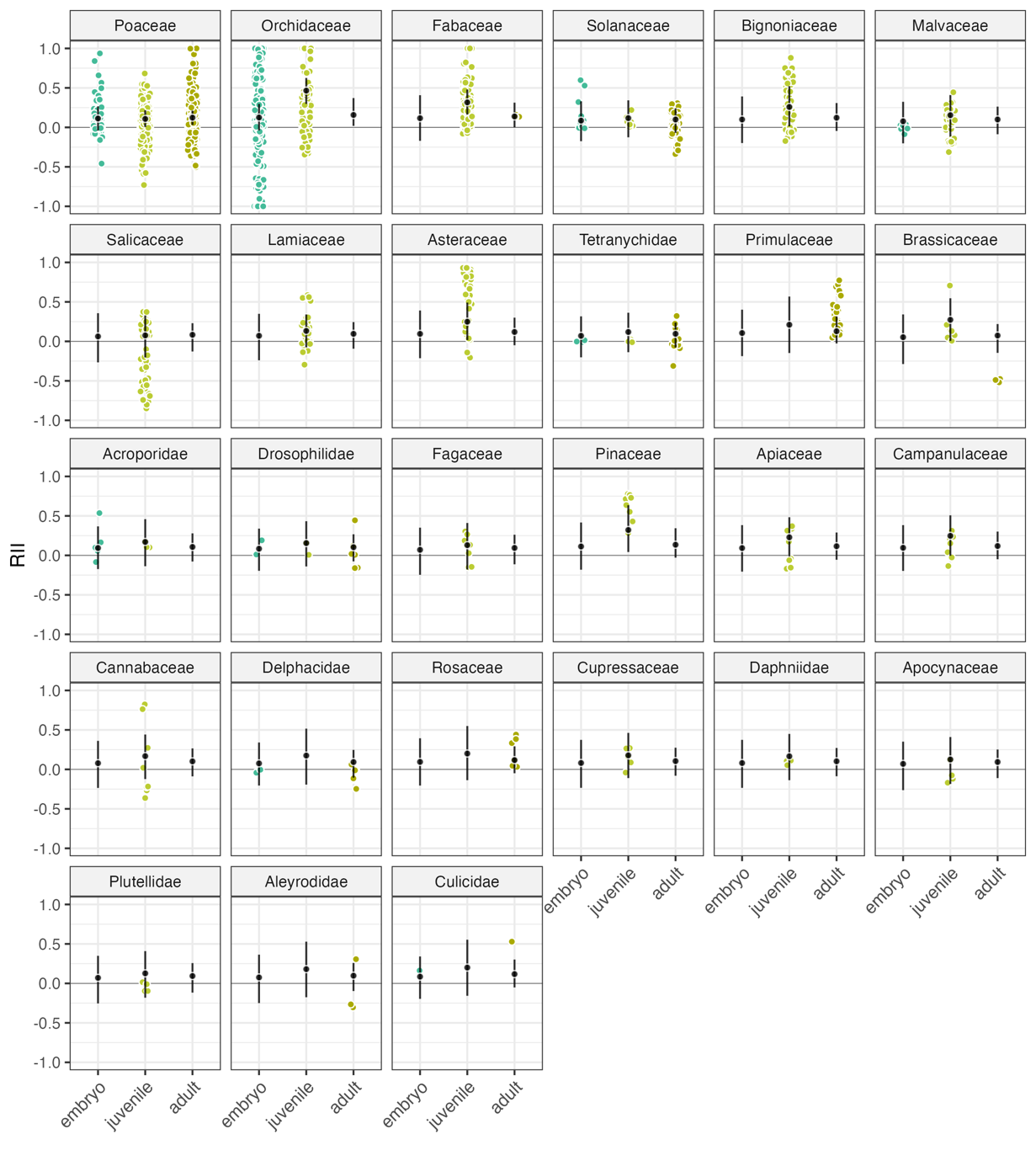

**Figure S12. Family specific meta-analytic mean estimates of microbial effects across host life stages.** Meta-analytic means (black) along with 95% CI are shown for each host life stage (embryo, juvenile, and adult), along with observed RII values across studies. Predictions averaging across taxonomic diversity are shown for each of 27 host families within the dataset. Panels for each family are presented in descending order of sample size.

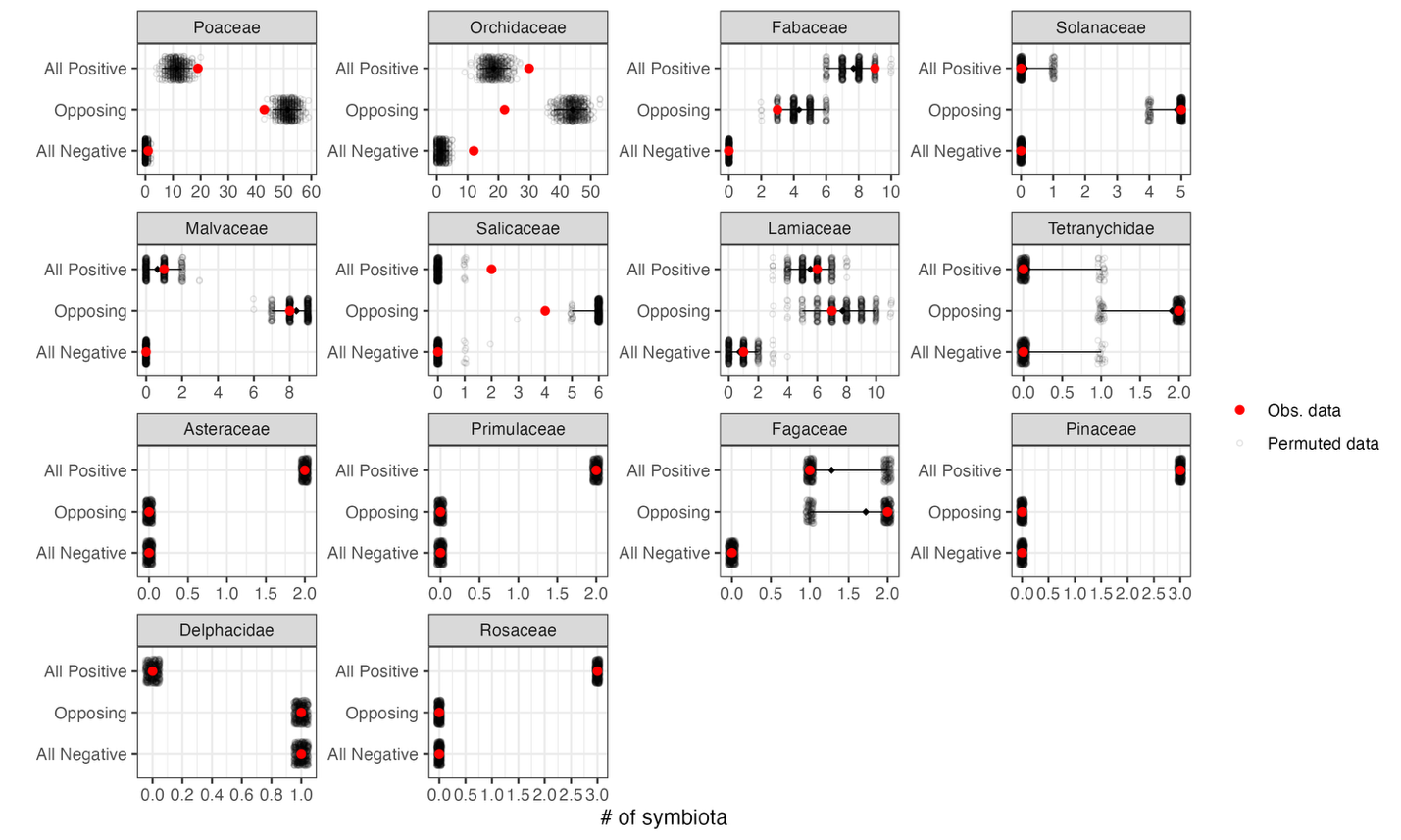

**Figure S13. Family specific frequency of opposing symbiotic effects.** Points represent the number of unique host-microbe symbiota with either all positive vital rate effects, at least one opposing vital rate effect, or all negative vital rate effects for each host family with more than two unique symbiota within the meta-analytic dataset (red) and across 500 randomly permuted datasets (black) for each family. Black diamonds represent the expected mean for each category from the permuted datasets along with error bars extending along the 95% CI. Panels for each family are presented in descending order of sample size, and note that families with few unique symbiota have fewer possible permutations.

**Identification of studies**

Records removed *before screening*:

Duplicate records removed (n = 0)

Records marked as ineligible by automation tools (n = 0)

Records removed for other reasons (n = 0)

Records identified from Web of Science (Feb. 21, 2024):

Databases (n = 1)

Records in search (n = 9606)

**Identification**

Records excluded:

- Does not evidently focus on host performance effects of symbiosis (n = 27)
- Review or meta-analysis (n = 9)

Records screened (Top 200 search results)

(n = 200)

Reports not accessible

(n = 8)

Reports sought for retrieval

(n = 164)

**Screening**

Reports excluded:

No explicit comparison of host performance with and without symbiont (n = 23)

Symbiont treatment combines multiple symbiont species in one treatment (n = 31)

Data inaccessible within report or uncollectable (i.e., no error bars, reported as grouped across multiple metrics) (n = 24)

etc.

Reports assessed for eligibility

(n = 156)

Total studies included in review

(n = 78)

**Included**

**Figure S14. PRISMA flow diagram of literature search and identification of included studies.** The first two hundred search results from a Web of Science were evaluated for inclusion, and then excluded for reasons including the lack of explicit comparison of host performance with and without a unique symbiont and for lack of data availability.

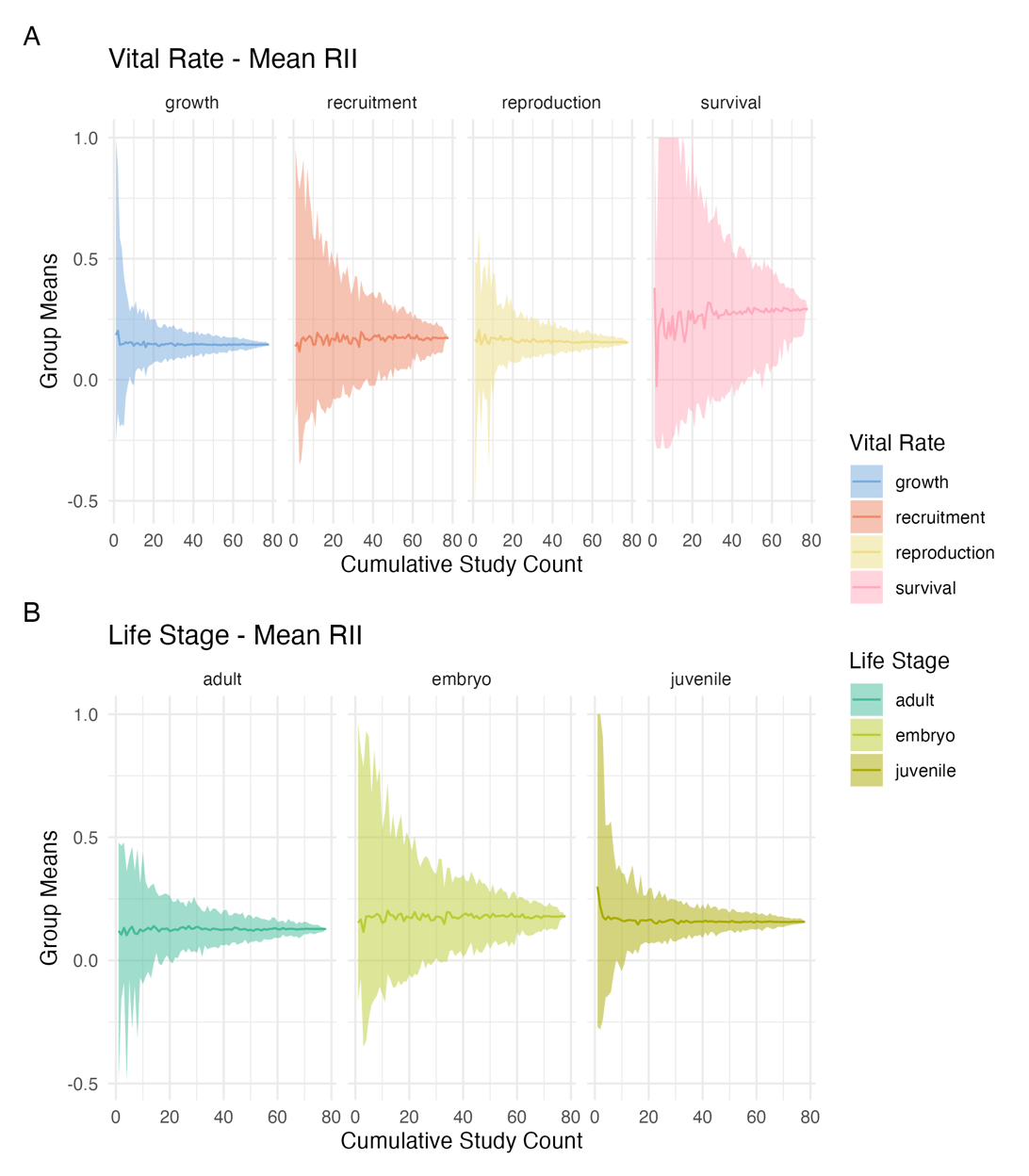

**Figure S15. Effect of meta-analysis dataset size on mean RII.** Group-wise means across (A) vital rates and (B) life stages calculated for randomly selected subsets of the data. Lines represent the mean and shaded areas show 95% confidence intervals calculated from 100 replicated subsets of studies randomly sampled from the full meta-analytic dataset (n = 78 studies). Each subset contains between 1 and 78 studies to evaluate how study inclusion influences estimated meta-analytic means.

**Supplemental Tables**

**Table S1. Categorization of Poaceae-associated symbiont partners**

| **Category** | **Symbiont Genera** | **# of associated host species** |
| --- | --- | --- |
| *Epichloë* fungi | *Epichloë* | 14 |
| Mycorrhizal fungi | *Acaulospora* | 1 |
|  | *Funneliformis* | 1 |
|  | *Gigaspora* | 1 |
|  | *Glomus* | 2 |
|  | *Rhizophagus* | 3 |
|  | *Serendipita* | 1 |
|  | *Xerocomus* | 1 |
|  | unk. mycorrhizal isolates | 2 |
| Bacteria | Achromobacter | 1 |
|  | Burkholderia | 1 |
|  | Methylobacterium | 1 |
|  | Microbacterium | 1 |
|  | Rhizobium | 1 |
|  | Streptomyces | 1 |
|  | unk. bacterial isolates | 2 |

**Table S2. List of included studies, measured fitness components and host and symbiont families.**

| **Authors** | **Year** | **Source Title** | **DOI** | **Vital Rates** | **Life Stages** | **Host Family** | **Symbiont Family** |
| --- | --- | --- | --- | --- | --- | --- | --- |
| Koch, AM; Croll, D; Sanders, IR | 2006 | ECOLOGY LETTERS | 10.1111/j.1461-0248.2005.00853.x | growth | juvenile | Lamiaceae, Poaceae | Glomeraceae |
| Rudgers, JA; Miller, TEX; Ziegler, SM; Craven, KD | 2012 | ECOLOGY | 10.1890/11-0689.1 | survival, growth, reproduction, recruitment | adult, embryo | Poaceae | Clavicipitaceae |
| Hosseini, A; Hosseini, M; Schausberger, P | 2022 | FRONTIERS IN PLANT SCIENCE | 10.3389/fpls.2021.783578 | growth, reproduction | adult | Rosaceae | Pseudomonadaceae, Rhodospirillaceae |
| Madhaiyan, M; Poonguzhali, S; Lee, HS; Hari, K; Sundaram, SP; Sa, TM | 2005 | BIOLOGY AND FERTILITY OF SOILS | 10.1007/s00374-005-0838-7 | recruitment, growth | embryo, juvenile | Poaceae | Methylobacteriaceae |
| Saeed, N; Battisti, A; Martinez-Sanudo, I; Mori, N | 2018 | BULLETIN OF INSECTOLOGY | NA | reproduction, recruitment, survival | adult, embryo, juvenile | Drosophilidae | Anaplasmataceae |
| Tian, F; Wang, JC; Bai, XX; Yang, YB; Huang, L; Liao, XF | 2023 | PLANT SIGNALING & BEHAVIOR | 10.1080/15592324.2023.2293405 | growth, recruitment, survival | juvenile, embryo | Orchidaceae | Mycorrhizal spp. |
| Karthikeyan, A; Arunprasad, T | 2021 | JOURNAL OF FORESTRY RESEARCH | 10.1007/s11676-019-01072-y | growth | juvenile | Fabaceae | Rhizobiaceae |
| Zhang, Y; Li, YY; Chen, XM; Guo, SX; Lee, YI | 2020 | BOTANICAL STUDIES | 10.1186/s40529-019-0278-6 | recruitment, survival | embryo, juvenile | Orchidaceae | Sebacinaceae, Tulasnellaceae |
| Wang, QX; Yan, N; Ji, DG; Li, SY; Hu, H | 2013 | BOTANICAL STUDIES | 10.1186/1999-3110-54-23 | growth | juvenile | Orchidaceae | Mycorrhizal spp. |
| Chen, XM; Dong, HL; Hu, KX; Sun, ZR; Chen, JA; Guo, SX | 2010 | J. OF PLANT GROWTH REGULATION | 10.1007/s00344-010-9139-y | growth, reproduction | juvenile | Orchidaceae | Mycorrhizal spp. |
| Hamilton, CE; Faeth, SH | 2005 | SYMBIOSIS | NA | growth | embryo | Poaceae | Clavicipitaceae |
| Wang, ZF; Li, CJ; White, J | 2020 | J. OF AGRONOMY AND CROP SCIENCE | 10.1111/jac.12366 | growth, reproduction, recruitment | juvenile, adult, embryo | Poaceae | Clavicipitaceae |
| Meng, PP; Chen, W; Feng, H; Zhang, SX; Wang, JH; Ma, WJ; Yang, GJ; Wang, CY | 2022 | NEW ZEALAND J. OF FORESTRY SCIENCE | 10.33494/nzifs522022x160x | growth | juvenile | Bignoniaceae | Glomeraceae |
| Otero, JT; Bayman, P; Ackerman, JD | 2005 | EVOLUTIONARY ECOLOGY | 10.1007/s10682-004-5441-0 | recruitment, growth | embryo | Orchidaceae | Ceratobasidiaceae, Meruliaceae |
| Niu, ZF; Yue, YS; Su, D; Ma, S; Hu, L; Hou, XC; Zhang, TT; Dong, D; Zhang, DP; Lu, CG; Fan, XF; Wu, HL | 2022 | BIOMASS & BIOENERGY | 10.1016/j.biombioe.2022.106360 | recruitment, growth | embryo, juvenile | Poaceae | Streptomycetaceae |
| Moore, JD; Carlisle, AE; Nelson, JA; McCulley, RL | 2019 | RESTORATION ECOLOGY | 10.1111/rec.12960 | growth, reproduction | adult | Poaceae | Clavicipitaceae |
| Li, HY; Smith, FA; Dickson, S; Holloway, RE; Smith, SE | 2008 | NEW PHYTOLOGIST | 10.1111/j.1469-8137.2008.02410.x | growth | juvenile | Poaceae | Glomeraceae, Gigasporaceae |
| Liu, S; Lv, DH; Lu, C; Xiao, YP; Wang, SQ; Zhou, W; Niu, JF; Wang, ZZ | 2022 | RHIZOSPHERE | 10.1016/j.rhisph.2022.100527 | recruitment, growth | embryo, juvenile | Orchidaceae | Tulasnellaceae |
| Liu, BH; Liu, XH; Liu, FC; Ma, HL; Ma, BY; Zhang, WX; Peng, L | 2019 | AMB EXPRESS | 10.1186/s13568-019-0865-7 | recruitment, growth | embryo, juvenile | Poaceae | Boletaceae |
| Rangjaroen, C; Rerkasem, B; Teaumroong, N; Noisangiam, R; Lumyong, S | 2015 | ANNALS OF MICROBIOLOGY | 10.1007/s13213-014-0857-4 | growth | juvenile | Poaceae | Mycorrhizal spp. |
| Hazzoumi, Z; Azaroual, SE; El Mernissi, N; Zaroual, Y; Duponnois, R; Bouizgarne, B; Kadmiri, IM | 2022 | FRONTIERS IN MICROBIOLOGY | 10.3389/fmicb.2022.881442 | growth | juvenile | Poaceae | Glomeraceae, Acaulosporaceae |
| Oien, DI; O'Neill, JP; Whigham, DF; McCormick, MK | 2008 | ANNALES BOTANICI FENNICI | 10.5735/085.045.0301 | recruitment, growth | embryo | Orchidaceae | Tulasnellaceae, Ceratobasidiaceae, |
| He, C; Wang, WQ; Hou, JL | 2019 | FRONTIERS IN MICROBIOLOGY | 10.3389/fmicb.2019.01364 | growth | juvenile | Fabaceae | Lophiostomataceae, Didymellaceae |
| Barazani, O; Benderoth, M; Groten, K; Kuhlemeier, C; Baldwin, IT | 2005 | OECOLOGIA | 10.1007/s00442-005-0193-2 | recruitment, growth, reproduction | embryo, juvenile, adult | Solanaceae | Mycorrhizal spp., Sebacinaceae |
| Wu, C; Wei, Q; Deng, J; Zhang, WY | 2019 | J. OF FORESTRY RESEARCH | 10.1007/s11676-018-0712-8 | growth | juvenile | Cupressaceae | Mycorrhizal spp. |
| Mala, B; Kuegkong, K; Sa-ngiaemsri, N; Nontachaiyapoom, S | 2017 | SOUTH AFRICAN JOURNAL OF BOTANY | 10.1016/j.sajb.2017.05.008 | recruitment, growth | embryo, juvenile | Orchidaceae | Tulasnellaceae |
| Shymanovich, T; Faeth, SH | 2019 | ECOLOGY AND EVOLUTION | 10.1002/ece3.5241 | growth | juvenile | Poaceae | Clavicipitaceae |
| Shao, SC; Luo, Y; Jacquemyn, H | 2020 | FRONTIERS IN PLANT SCIENCE | 10.3389/fpls.2020.571426 | recruitment | embryo | Orchidaceae | Mycorrhizal spp., Tulasnellaceae |
| Chen, W; Meng, PP; Feng, H; Wang, CY | 2020 | FORESTS | 10.3390/f11101117 | growth | juvenile | Bignoniaceae | Glomeraceae |
| Chung, YA; Miller, TEX; Rudgers, JA | 2015 | JOURNAL OF ECOLOGY | 10.1111/1365-2745.12406 | survival, growth, reproduction, recruitment | adult, juvenile, embryo | Poaceae | Clavicipitaceae |
| Decruse, SW; Neethu, RS; Pradeep, NS | 2018 | S. AFRICAN J. OF BOTANY | 10.1016/j.sajb.2018.04.002 | recruitment, survival | embryo, juvenile | Orchidaceae | Mycorrhizal spp. |
| Saxena, J; Minaxi; Jha, A | 2014 | CLEAN-SOIL AIR WATER | 10.1002/clen.201300492 | recruitment, growth, reproduction | embryo, juvenile, adult | Poaceae | Glomeraceae, Burkholderiaceae |
| Jansa, J; Smilauer, P; Borovicka, J; Hrselov√°, H; Forczek, ST; Sl√°mov√°, K; Rezanka, T; Rozmos, M; Bukovsk√°, P; Gryndler, M | 2020 | MYCORRHIZA | 10.1007/s00572-020-00937-z | growth | juvenile | Poaceae | Glomeraceae |
| Bao, GS; Song, ML; Wang, YQ; Saikkonen, K; Wang, HS | 2019 | SYMBIOSIS | 10.1007/s13199-019-00636-0 | recruitment, growth | embryo | Poaceae | Clavicipitaceae |
| Soares, MA; Li, HY; Kowalski, KP; Bergen, M; Torres, MS; White, JF | 2016 | MICROBIAL ECOLOGY | 10.1007/s00248-016-0793-x | growth | juvenile | Poaceae | Alcaligenaceae, Microbacteriaceae |
| Meier, AR; Hunter, MD | 2018 | FRONT. IN ECO. AND EVO. | 10.3389/fevo.2018.00033 | growth | juvenile | Apocynaceae | Glomeraceae |
| Langill, T; Jorissen, LP; Ole√±ska, E; W√≥jcik, M; Vangronsveld, J; Thijs, S | 2023 | PLANTS-BASEL | 10.3390/plants12030643 | growth, recruitment, reproduction | juvenile, adult | Brassicaceae | Sphingomonadaceae |
| Zhang, X; Ohtsuki, H; Makino, W; Kato, Y; Watanabe, H; Urabe, J | 2021 | ECOLOGICAL RESEARCH | 10.1111/1440-1703.12194 | growth | juvenile | Daphniidae | Comamonadaceae, Enterobacteriaceae |
| Hoffmann, D; Vierheilig, H; Peneder, S; Schausberger, P | 2011 | ECOLOGICAL ENTOMOLOGY | 10.1111/j.1365-2311.2011.01298.x | growth, reproduction | juvenile, adult | Fabaceae | Glomeraceae |
| Dupin, SE; Geurts, R; Kiers, ET | 2020 | FRONTIERS IN PLANT SCIENCE | 10.3389/fpls.2019.01779 | growth | juvenile | Cannabaceae | Bradyrhizobiaceae |
| Gibert, A; Hazard, L | 2011 | JOURNAL OF PLANT ECOLOGY | 10.1093/jpe/rtr009 | growth, recruitment | juvenile, embryo | Poaceae | Clavicipitaceae |
| Jiang, XL; Zhao, ZY; Jacquemyn, H; Ding, G; Ding, WL; Xing, XK | 2022 | GLOBAL ECOLOGY AND CONSERVATION | 10.1016/j.gecco.2022.e02235 | recruitment | embryo | Orchidaceae | Ceratobasidiaceae |
| Gundel, PE; Zabalgogeazcoa, I; de Aldana, BRV | 2011 | CROP & PASTURE SCIENCE | 10.1071/CP11300 | recruitment | adult | Poaceae | Clavicipitaceae |
| Emery, SM; Rudgers, JA | 2013 | OECOLOGIA | 10.1007/s00442-013-2705-9 | survival, growth | juvenile | Poaceae | Clavicipitaceae |
| Duponnois, R; Founoune, H; Lesueur, D | 2002 | GEODERMA | 10.1016/S0016-7061(02)00144-1 | growth | juvenile | Fabaceae | Sclerodermataceae, Bradyrhizobiaceae |
| Gundel, PE; Helander, M; Casas, C; Hamilton, CE; Faeth, SH; Saikkonen, K | 2013 | FUNGAL DIVERSITY | 10.1007/s13225-012-0173-x | growth, reproduction | adult | Poaceae | Clavicipitaceae |
| Liu, T; Wang, CY; Chen, H; Fang, FR; Zhu, XQ; Tang, M | 2014 | ACTA PHYSIOLOGIAE PLANTARUM | 10.1007/s11738-013-1465-9 | growth | juvenile | Salicaceae | Glomeraceae |
| Larcher, M; Rapior, S; Cleyet-Marel, JC | 2008 | ACTA BOTANICA GALLICA | 10.1080/12538078.2008.10516116 | growth | juvenile | Brassicaceae | Phyllobacteriaceae |
| Xie, RR; Sun, JT; Xue, XF; Hong, XY | 2016 | SYSTEMATIC AND APPLIED ACAROLOGY | 10.11158/saa.21.9.1 | reproduction, recruitment, survival | adult, embryo, juvenile | Tetranychidae | Anaplasmataceae, Flammeovirgaceae |
| Hazraty-Kari, S; Masaya, M; Kawachi, M; Harii, S | 2022 | J OF EXPERIMENTAL ZOOLOGY | 10.1002/jez.2589 | survival, growth, recruitment | embryo, juvenile | Acroporidae | Suessiaceae |
| Sudov√°, R; Rydlov√°, J; M√ºnzbergova, Z; Suda, J | 2010 | AMERICAN JOURNAL OF BOTANY | 10.3732/ajb.1000114 | growth | juvenile | Asteraceae, Campanulaceae, Apiaceae | Glomeraceae |
| Gibert, A; Hazard, L | 2013 | JOURNAL OF ECOLOGY | 10.1111/1365-2745.12073 | growth, reproduction | adult | Poaceae | Clavicipitaceae |
| Wang, L; Chen, X; Wang, S; Du, YQ; Zhang, D; Tang, ZH | 2022 | EUROPEAN J. OF SOIL BIOLOGY | 10.1016/j.ejsobi.2022.103429 | growth, reproduction, recruitment | adult | Solanaceae | Glomeraceae |
| Ilyas, F; Ali, MA; Modhish, A; Ahmed, N; Hussain, S; Bilal, M; Arshad, M; Danish, S; Ghoneim, AM; Ilyas, A; Akram, A; Fahad, S; Ansari, MJ; Datta, R | 2023 | CROP & PASTURE SCIENCE | 10.1071/CP21042 | recruitment, growth, reproduction | embryo, adult | Poaceae | Clavicipitaceae |
| Haidar, B; Ferdous, M; Fatema, B; Ferdous, AS; Islam, MR; Khan, H | 2018 | MICROBIOLOGICAL RESEARCH | 10.1016/j.micres.2018.01.008 | recruitment, growth | embryo, juvenile | Malvaceae | Micrococcaceae, Staphylococcaceae, Burkholderiaceae, Brevibacteriaceae, Bacillaceae |
| Cheplick, GP | 2011 | AMERICAN JOURNAL OF BOTANY | 10.3732/ajb.1000226 | growth, reproduction | juvenile, adult | Poaceae | Clavicipitaceae |
| Somerville, J; Zhou, LQ; Raymond, B | 2019 | INSECTS | 10.3390/insects10040089 | survival, growth | juvenile | Plutellidae | Enterobacteriaceae |
| Zong, K; Huang, J; Nara, K; Chen, YH; Shen, ZG; Lian, CL | 2015 | JOURNAL OF FOREST RESEARCH | 10.1007/s10310-015-0506-1 | survival, growth | juvenile | Pinaceae, Fagaceae | Sclerodermataceae, Gloniaceae, Hydnangiaceae |
| Gao, YY; Peng, SJ; Hang, Y; Xie, GF; Ji, N; Zhang, MS | 2022 | SCIENTIA HORTICULTURAE | 10.1016/j.scienta.2021.110724 | recruitment | embryo | Orchidaceae | Psathyrellaceae |
| Li, TP; Zhou, CY; Zha, SS; Gong, JT; Xi, ZY; Hoffmann, AA; Hong, XY | 2020 | APPLIED AND ENV. MICROBIOLOGY | 10.1128/AEM.02509-19 | reproduction, recruitment, survival | adult, embryo | Delphacidae | Flammeovirgaceae, Anaplasmataceae |
| Campo, S; Mart√≠n-Cardoso, H; Oliv√©, M; Pla, E; Catala-Forner, M; Mart√≠nez-Eixarch, M; San Segundo, B | 2020 | RICE | 10.1186/s12284-020-00402-7 | growth, reproduction | juvenile, adult, embryo | Poaceae | Glomeraceae |
| Xie, RR; Chen, XL; Hong, XY | 2011 | APPLIED ENTOMOLOGY AND ZOOLOGY | 10.1007/s13355-010-0014-x | reproduction, recruitment, growth, survival | adult, embryo, juvenile | Tetranychidae | Anaplasmataceae |
| Rooney, DC; Prosser, JI; Bending, GD; Baggs, EM; Killham, K; Hodge, A | 2011 | BIOMASS & BIOENERGY | 10.1016/j.biombioe.2011.08.015 | growth | juvenile | Salicaceae | Glomeraceae, Gigasporaceae |
| Lahrizi, Y; Oukaltouma, K; Mouradi, M; Farissi, M; Qaddoury, A; Bouizgaren, A; Ghoulam, C | 2021 | APPLIED ECO. AND ENV. RESEARCH | 10.15666/aeer/1901_563580 | growth | juvenile | Fabaceae | Rhizobiaceae |
| Granada, CE; Arruda, L; Lisboa, BB; Passaglia, LMP; Vargas, LK | 2014 | BIOLOGY AND FERTILITY OF SOILS | 10.1007/s00374-013-0840-4 | growth, recruitment | juvenile, embryo | Fabaceae, Poaceae | Rhizobiaceae |
| Sudov√°, R; P√°nkov√°, H; Rydlov√°, J; M√ºnzbergov√°, Z; Suda, J | 2014 | AMERICAN JOURNAL OF BOTANY | 10.3732/ajb.1300262 | growth | juvenile | Asteraceae | Glomeraceae |
| Rydlov√°, J; Sykorov√°, Z; Slav√≠kov√°, R; Turis, P | 2015 | MYCORRHIZA | 10.1007/s00572-015-0634-7 | growth, reproduction | adult | Primulaceae | Mycorrhizal spp. |
| Dobson, SL; Rattanadechakul, W; Marsland, EJ | 2004 | HEREDITY | 10.1038/sj.hdy.6800458 | recruitment, reproduction | embryo, adult | Culicidae | Anaplasmataceae |
| Faeth, SH; Hayes, CJ; Gardner, DR | 2010 | MICROBIAL ECOLOGY | 10.1007/s00248-010-9643-4 | survival, growth, reproduction | adult | Poaceae | Clavicipitaceae |
| Msaddak, A; Quinones, MA; Mars, M; Pueyo, JJ | 2023 | PLANTS-BASEL | 10.3390/plants12244109 | growth | juvenile | Fabaceae | Rhizobia spp. |
| Iannone, LJ; Pinget, AD; Nagabhyru, P; Schardl, CL; De Battista, JP | 2012 | GRASS AND FORAGE SCIENCE | 10.1111/j.1365-2494.2012.00855.x | growth, reproduction | juvenile, adult | Poaceae | Clavicipitaceae |
| Verlinden, MS; Ven, A; Verbruggen, E; Janssens, IA; Wallander, H; Vicca, S | 2018 | ECOLOGY | 10.1002/ecy.2502 | growth, reproduction | adult | Poaceae | Glomeraceae |
| Zi, XM; Sheng, CL; Goodale, UM; Shao, SC; Gao, JY | 2014 | MYCORRHIZA | 10.1007/s00572-014-0565-8 | recruitment, growth | embryo, juvenile | Orchidaceae | Tulasnellaceae, Hypocreaceae |
| Aewsakul, N; Maneesorn, D; Serivichyaswat, P; Taluengjit, A; Nontachaiyapoom, S | 2013 | SCIENTIA HORTICULTURAE | 10.1016/j.scienta.2013.05.034 | recruitment, growth | embryo, juvenile | Orchidaceae | Tulasnellaceae |
| Wu, YH; Wang, H; Liu, M; Li, B; Chen, X; Ma, YT; Yan, ZY | 2021 | FRONTIERS IN PLANT SCIENCE | 10.3389/fpls.2021.617892 | growth | juvenile | Lamiaceae | Glomeraceae, Ambisporaceae, Acaulosporaceae |
| Tian, F; Liao, XF; Wang, LH; Bai, XX; Yang, YB; Luo, ZQ; Yan, FX | 2022 | PLANT SIGNALING & BEHAVIOR | 10.1080/15592324.2021.2005882 | growth | juvenile | Orchidaceae | Tulasnellaceae |
| Ren, FR; Sun, X; Wang, TY; Yan, JY; Yao, YL; Li, CQ; Luan, JB | 2021 | ISME JOURNAL | 10.1038/s41396-020-00877-8 | survival, reproduction | adult | Aleyrodidae | Halomonadaceae |
| Huang, H; Zi, XM; Lin, H; Gao, JY | 2018 | JOURNAL OF MICROBIOLOGY | 10.1007/s12275-018-7225-1 | recruitment, growth | embryo, juvenile | Orchidaceae | Tulasnellaceae |
